# Phage-encoded homing endonucleases attenuate bacterial immunity

**DOI:** 10.1101/2025.09.07.674715

**Authors:** Tridib Mahata, Katarzyna Kanarek, Rameshkumar Marimuthu Ragavan, Anushree Haldar, Guy Shur, Ruba Yehia, David Burstein, Yoni Haitin, Udi Qimron, Dor Salomon

## Abstract

The arms race between bacteria and bacteriophages (phages) gave rise to multiple layers of antagonistic mechanisms, many of which remain unexplored. Here, we investigated the anti-phage defense system GAPS4 and showed that it is a non-selective DNase triggered by sensing DNA breaks. We further demonstrated that this activation mechanism renders GAPS4 a double-edged sword, sensitizing bacteria to various forms of antibacterial antagonism. Using comparative genomics, we found that phage-encoded homing endonucleases, long considered selfish mobile genetic elements, enhance phage fitness by attenuating GAPS4-mediated immunity. Our findings shed light on the evolutionary advantage provided by these ubiquitous mobile elements to their host phages, and on the intricate evolutionary cross-talk between bacteria and their predators.

## Introduction

Bacteriophages (phages) prey on bacteria, leading to an evolutionary arms race in which bacteria evolve anti-phage defense mechanisms(*1*) and phages evolve countermeasures to antagonize them(*2*). Anti-phage defense systems can recognize a conserved phage component, such as a phage protein or the incoming phage DNA, or a perturbation of a cellular process to mount either a restrictive or abortive response(*3*). In the former, the bacterium specifically targets and restricts the incoming phage, whereas in the latter, it targets itself, often activating a suicide program to prevent phage proliferation(*4*).

In less than a decade, >200 types of anti-phage defense systems have been described(*5, 6*), as well as several phage-encoded anti-defense mechanisms(*2*). We recently reported an anti-phage defense system named GAPS4, discovered in a mobile genetic element called the GMT island(*7*). We showed that GAPS4 from *Vibrio parahaemolyticus* MAVP-R (GAPS4^MAVP-R^) protects against various coliphages when ectopically expressed in an *E. coli* surrogate strain. Similar to most known anti-phage defense systems, the trigger, target, mechanism of action, and structure of GAPS4 remain unknown.

Here, we reveal that GAPS4 is a heterotetramer DNase that targets both phage and bacterial DNA upon sensing linear DNA during phage infection. Notably, sensing linear DNA as a trigger renders GAPS4 a double-edged sword, hyper-sensitizing its host to antibacterial antagonism that induces DNA breaks. Furthermore, we describe phage-encoded homing endonucleases as anti-defense systems that attenuate the bacterial immune response mediated by GAPS4.

## Results

### GAPS4 represents a widespread family of anti-phage defense systems

GAPS4^MAVP-R^ comprises two genes encoding GAPS4a and GAPS4b (Fig. 1a; WP_055466293.1 and WP_055466294.1). To determine whether both genes are required for anti-phage protection, we examined the ability of each gene to mount a defense against lytic phage λ attacks. We found that neither GAPS4a nor GAPS4b protects *E. coli* without its counterpart, indicating that both are essential components of the defense system (Fig. 1b).

**Fig. 1.**
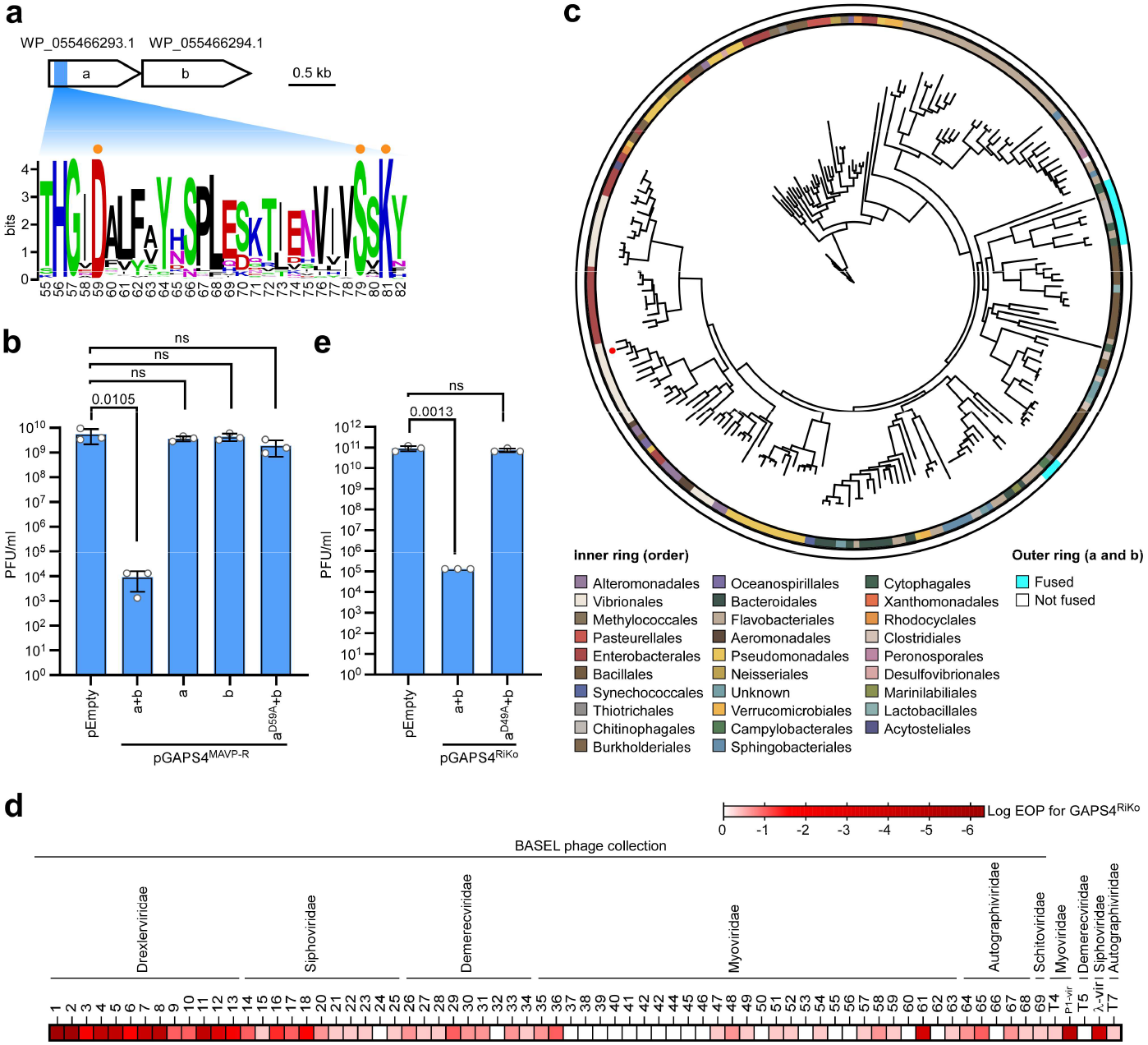
GAPS4 is a widespread family of anti-phage defense systems with a PD-(D/E)xK phosphodiesterase active domain. **(a)** Schematic representation of the GAPS4 system. The conservation logo of the predicted PD-(D/E)xK phosphodiesterase domain active site found in GAPS4 homologs (denoted by orange circles) is shown below. The residue numbers correspond to the positions in GAPS4a^MAVP-R^ (WP_055466293.1). **(b** and **e)** Plaque-forming units (Pfus) of λ-vir phage upon infection of *E. coli* strains containing an empty plasmid (pEmpty) or a plasmid for the arabinose-inducible expression of the indicated GAPS4^MAVP-R^ version or the expression of the indicated GAPS4^RiKo^ from its native promoter (e). Statistical significance between samples was calculated using One-way ANOVA with Dunnett multiple comparisons correction; ns, no significant difference (*P* > 0.05). Data are shown as mean ± SD of 3 independent experiments. **(c)** Phylogenetic distribution of GAPS4 homologs. The bacterial order is denoted by color. The outer ring denotes GAPS4 homologs naturally fused as one gene. The evolutionary history was inferred using RAxML on a concatenated alignment of GAPS4a and GAPS4b along with fused GAPS4a-b proteins. GAPS4^MAVP-R^ is denoted with a red circle. **(d)** The efficiency of plating (EOP; shown as the log difference in plaque numbers) was determined for *E. coli* harboring a low-copy plasmid encoding GAPS4^RiKo^ with its native promoter, when challenged with 73 coliphages, compared to *E. coli* containing an empty plasmid. The data shown are the average of 3 independent experiments.

Computational analyses revealed 1,639 non-redundant GAPS4 homologs (comprising GAPS4a and GAPS4b in tandem or fused as a single gene) in 59 different bacterial orders, including gram-negative and gram-positive bacteria inhabiting diverse ecological niches (Fig. 1c, Dataset S1, and File S1). To determine whether GAPS4 homologs also function as anti-phage defense systems, we subcloned a GAPS4 homolog from *E. coli* strain RiKo 2299/09 (GAPS4^RiKo^; WP_028132563.1 and WP_028132564.1; 30.63% identity with 88% query coverage and 25% identity with 97% query coverage compared to GAPS4a^MAVP-R^ and GAPS4b^MAVP-R^, respectively), together with its native promoter, into a low-copy plasmid. We observed that a phage-sensitive, surrogate *E. coli* strain harboring this plasmid became resistant to attacks by multiple phages compared to *E. coli* containing an empty plasmid, as evidenced by a reduction of up to six orders of magnitude in the number of visible plaques that developed on a lawn of bacteria (i.e., the efficiency of plating; EOP) (Fig. 1d).

Amino acid conservation analyses of GAPS4a and GAPS4b homologs (Fig. S1 and Fig. S2) implied that GAPS4a belongs to a divergent family of PD-(D/E)xK phosphodiesterases with the non-canonical conserved motif HGxD-SxK(*8, 9*) at its N-terminus (Fig. 1a). In agreement with this prediction, substitution of the conserved aspartic acid in position 59 with alanine abrogated GAPS4^MAVP-R^-mediated protection against lytic phage λ attacks (Fig. 1b). This conserved aspartic acid was also required for GAPS4^RiKo^-mediated protection (Fig. 1e). Collectively, our results indicate that GAPS4 is a widespread family of anti-phage defense systems with a predicted PD-(D/E)xK phosphodiesterase active domain.

### GAPS4 is a heterotetrameric complex

The results above indicated that both GAPS4 components are essential for its activity. Therefore, we hypothesized that GAPS4a and GAPS4b form a complex. Indeed, when we purified GAPS4a using size-exclusion chromatography (SEC) from an *E. coli* strain expressing the GAPS4^MAVP-R^ operon (GAPS4a and GAPS4b), GAPS4b co-purified with it, supporting our hypothesis (Fig. S3a). Mass spectrometry analysis of the peak fractions confirmed that GAPS4a and GAPS4b are the two prevalent proteins in the eluate (data not shown). Notably, the SEC elution profile suggested a possible higher oligomeric state of the GAPS4 complex. To evaluate the stoichiometry and absolute molar mass of the purified GAPS4 complex, we used size-exclusion chromatography multi-angle light scattering (SEC-MALS). As shown in Fig. S3b, we observed a mono-dispersed complex with a calculated mass of 164.3 ± 11.5 kDa, corresponding with the predicted mass of a heterotetramer comprising two GAPS4a molecules and two GAPS4b molecules (160.8 kDa).

In agreement with the SEC-MALS results, AlphaFold3 analysis predicts, with high confidence, that GAPS4a and GAPS4b of GAPS4^MAVP-R^ form a heterotetramer with a 2:2 stoichiometry (Fig. S3c). In this model, the two GAPS4b subunits form a ring-like structure, and the GAPS4a subunits are docked onto α-helices-comprising globular domains in the GAPS4b subunits. The predicted active sites of the GAPS4a subunits, comprising the conserved HGxD-SxK motif, face outwards to opposite sides of the GAPS4b ring (Fig. S3c and File S2). A similar complex is predicted for GAPS4^RiKo^ (Fig. S4 and File S3).

### GAPS4 degrades phage and bacterial DNA

Our analyses indicated that GAPS4 protects *E. coli* against several lytic phages, including λ-vir, an obligatory lytic variant of the λ phage (Fig. 1)(*7*). To determine whether GAPS4 can also protect bacteria against a lysogenic phage, we challenged *E. coli* cells harboring either a plasmid-borne GAPS4^RiKo^ or its inactive mutant GAPS4^RiKo(D49A)^ with λ phage harboring a kanamycin-resistance cassette, under lysogenization-favoring conditions. We obtained ∼100-1000-fold fewer lysogens in the presence of the wild-type GAPS4, whether bacteria were challenged with the phage at a multiplicity of infection (MOI) of 0.1 or 5 (Fig. 2a). These results confirm that GAPS4 provides protection against lytic and lysogenic phage attacks.

**Fig. 2.**
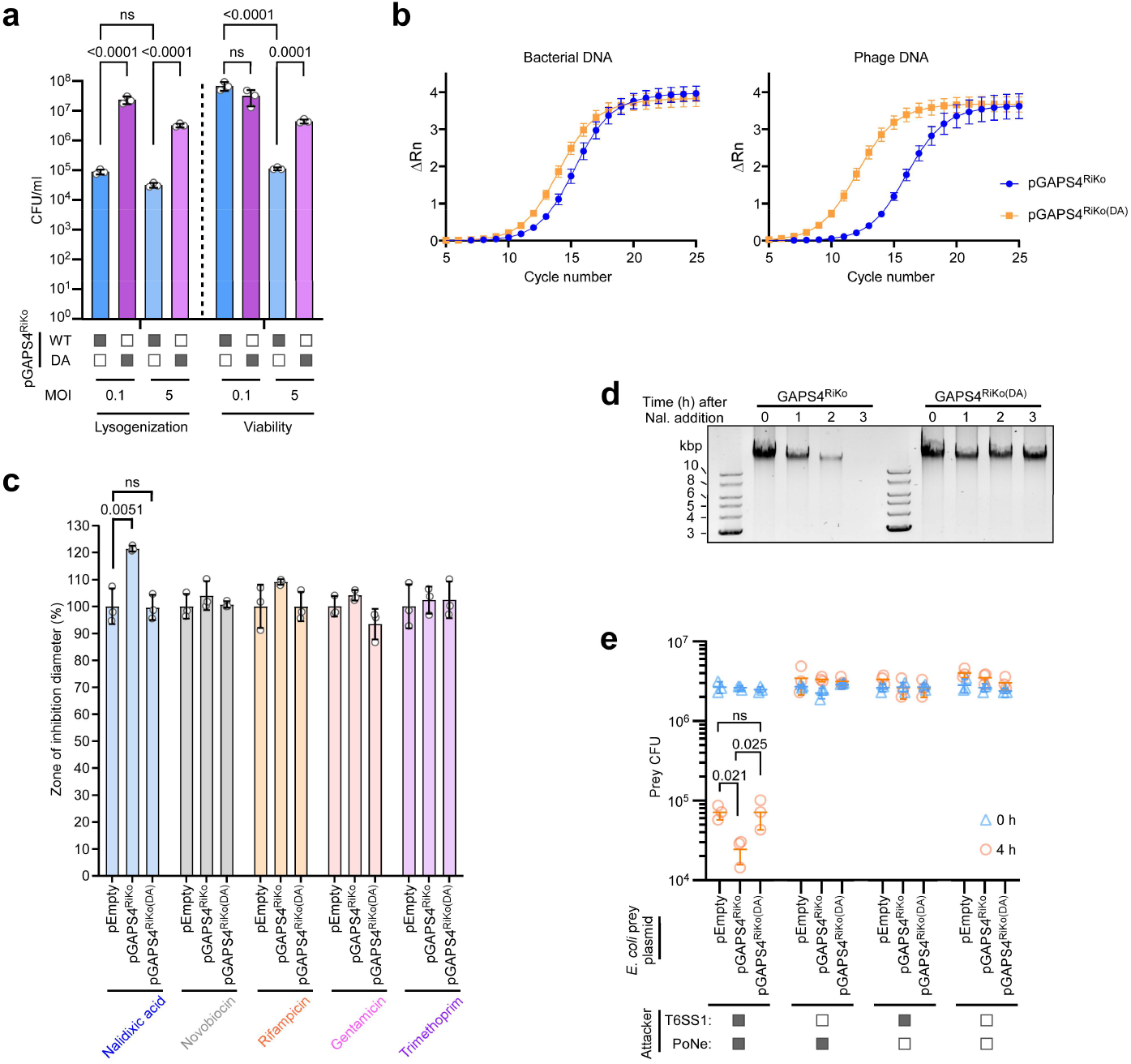
GAPS4 degrades DNA upon sensing linear DNA. **(a)** Lysogenization and viability of *E. coli* strains containing the indicated GAPS4^RiKo^ upon challenge with λ phage under lysogenization-favourable conditions. MOI, multiplicity of infection; WT, wild-type; DA, D59A mutant. Statistical significance between samples was calculated using Two-way ANOVA with Tukey multiple comparisons correction on log-transformed data. Data are shown as mean ± SD of 3 independent samples. The experiment was repeated three times with similar results. Results from a representative experiment are shown. **(b)** Bacterial and phage chromosomal DNA stability as determined by quantitative RT-PCR, 90 minutes after *E. coli* strains harboring a plasmid expressing GAPS4^RiKo^ (pGAPS4^RiKo^) or its catalytic mutant (pGAPS4^RiKo(DA)^) were challenged with λ phage under lysogenization-favourable conditions at MOI = 1. Data are shown as mean ± SD of 3 independent experiments, each with 3 technical repeats. ΔRn, normalized change in qPCR fluorescence signal. Sensitivity of *E. coli* strains harboring an empty plasmid (pEmpty) or a plasmid expressing GAPS4^RiKo^ (pGAPS4^RiKo^) or its catalytic mutant (pGAPS4^RiKo(DA)^) to the indicated antibiotics, measured as the zone of inhibition diameter around a filter disc containing the antibiotics. Statistical significance between samples was calculated using an unpaired, two-tailed Student *t*-test. Data are shown as mean ± SD of 3 independent experiments. **(d)** Genomic DNA stability after the addition of nalidixic acid (Nal.; 5 µg ml^−1^) to the media of *E. coli* cultures containing the indicated GAPS4^RiKo^ on a plasmid. The experiment was repeated three times with similar results. Results from a representative experiment are shown. **(e)** Viability counts (colony forming units; CFU) of *E. coli* prey strains containing the indicated plasmids before (0 h) and after (4 h) co-incubation with *Vibrio parahaemolyticus* attacker strains lacking their endogenous T6SS antibacterial effectors, with either a functional (full square) or inactive (empty square) T6SS1, containing either an empty plasmid or a plasmid for expression of the DNase T6SS1 effector PoNe. Statistical significance between samples at the 4 h time point was calculated using One-way ANOVA with Tukey multiple comparisons correction on log-transformed data. Data are shown as mean ± SD of 3 independent samples. The experiment was repeated three times with similar results. In a, c, and e: ns, no significant difference (*P* > 0.05).

GAPS4’s ability to protect against a lysogenic phage prompted us to investigate whether it restricts the incoming phage or induces bacterial cell suicide upon invasion. Although we did not detect a significant change in bacterial viability upon phage challenge at MOI 0.1 (when <10% of bacteria are exposed to phage attack), we noted a significant reduction in viability (∼40-fold) at MOI 5 (when most bacteria are exposed to phage attack) in the presence of GAPS4^RiKo^ (Fig. 2a). If the system merely provided protection against the phage rather than induced cell suicide, we would expect to observe minimal loss of viability. Therefore, these results suggested that GAPS4 induces cell suicide upon phage attack under lysogenization-favoring conditions.

Many members of the PD-(D/E)xK phosphodiesterase superfamily cleave DNA(*10*). Therefore, we hypothesized that GAPS4 is a DNase that induces cell suicide by degrading the bacterial genomic DNA upon phage attack. Indeed, when we monitored the levels of bacterial genomic DNA using quantitative real-time PCR, we found that upon challenge with λ phage under lysogenization-favoring conditions at MOI 1, the presence of GAPS4^RiKo^ led to a decrease in DNA compared to its inactive mutant form (manifested as delayed signal accumulation) (Fig. 2b). We also observed genomic DNA degradation upon infection of *E. coli* harboring GAPS4^MAVP-R^ with the lytic phages λ-vir and BASEL_01 (Fig. S5a). Remarkably, we found that GAPS4^RiKo^ also mediated the apparent degradation of the incoming λ phage DNA, suggesting that GAPS4 degrades both phage and host DNA upon infection (Fig. 2b).

### GAPS4 is a sentinel of DNA integrity

Following these results, we sought to identify the trigger that induces GAPS4-mediated DNA degradation. Despite repeated attempts, we were unable to isolate escape phages that bypass GAPS4-mediated defense, leading us to hypothesize that instead of sensing a phage protein, GAPS4 senses the injection of phage DNA or a perturbation to the host cell that occurs during the infection. Because we did not detect GAPS4-mediated cell death upon the presence of lysogenized λ phage DNA in the bacterial genome (Fig. S6a) or upon plasmid DNA injection by a transducing phage T7 particle (Fig. S6b), we reasoned that GAPS4 is not triggered by the phage-mediated injection process but rather senses a host cell perturbation.

To mimic possible phage-induced perturbations to host cellular processes, we employed various classes of antibiotics that inhibit DNA synthesis (Trimethoprim), replication (Nalidixic acid and Novobiocin), transcription (Rifampicin), or translation (Gentamicin). When we determined the susceptibility of *E. coli* strains harboring wild-type or mutant GAPS4^RiKo^ to the various antibacterial compounds, the wild-type GAPS4^RiKo^ specifically sensitized bacteria to nalidixic acid, a gyrase inhibitor predicted to induce DNA breaks, as evidenced by a larger inhibition zone in a disc diffusion assay (Fig. 2c). Sensitivity to nalidixic acid, but not to other antibiotic compounds not predicted to induce DNA breaks, was also observed with GAPS4^MAVP-R^ (Fig. S5b). Importantly, exposure to nalidixic acid in the presence of GAPS4^RiKo^ or GAPS4^MAVP-R^ led to degradation of the bacterial genomic DNA (Fig. 2d, Fig. S5a, and Fig. S5c). Collectively, these results demonstrate that nalidixic acid specifically triggers GAPS4 activity, leading to DNA degradation.

### GAPS4 sensitizes its host to antibacterial mechanisms that induce DNA breaks

Various mechanisms of antibacterial antagonism can induce DNA breaks. For example, Type VI secretion systems (T6SSs) are offensive antibacterial weapons that deliver toxic proteins, called effectors, directly into neighboring target cells(*11*). Because many T6SS effectors are DNases(*12*), we hypothesized that if GAPS4 is triggered by DNA breaks, then it can hyper-sensitize recipient bacteria to T6SS-mediated DNase attacks. To test this, we monitored the viability of *E. coli* prey cells harboring wild-type or mutant forms of GAPS4^RiKo^ after 4 hours of co-incubation with attacking bacteria that specifically deliver a single T6SS DNase effector, PoNe(*13*). In agreement with our hypothesis, prey cells harboring the wild-type GAPS4^RiKo^ were more susceptible to PoNe-mediated toxicity, as evidenced by ∼3-fold decrease in viability compared to prey cells harboring GAPS4^RiKo(D59A)^ or an empty plasmid (Fig. 2e). These results further support the conclusion that GAPS4 senses DNA breaks. Furthermore, they suggest that GAPS4 is a double-edged sword that protects its host from invading phages while sensitizing it to certain forms of antibacterial antagonism.

### A phage-borne homing endonuclease attenuates GAPS4-mediated protection

While inspecting the protection provided by GAPS4^RiKo^ against various phages, we noted different levels of protection (i.e., EOP) against phages belonging to the same family and having similar genomic composition (Fig. 1d). Phages BASEL_21 – BASEL_25 are members of the *Siphoviridae* family, yet only BASEL_24 infection is not hindered by GAPS4^RiKo^, compared to other family members that are mildly inhibited. Therefore, we hypothesized that the BASEL_24 phage harbors a phage anti-defense system.

Comparative genomics analysis of the BASEL_21 – BASEL_25 genomes revealed one gene that is unique to BASEL_24: *bas24_0090*, encoding a putative homing endonuclease with an HNH DNase domain (similar to I-HmuI(*14*); 99.94% probability according to HHpred(*15*) analysis) (Fig. 3a). Homing endonucleases, which are ubiquitous in phage and bacterial genomes, are considered selfish elements that can insert their encoding gene into specific DNA sites(*16*). They are modular proteins comprising a DNA-binding domain and a nuclease domain(*17*), and they include several families that have been proposed to undergo domain shuffling(*18*). Although thought to contribute to phage evolution(*18*), the advantage that homing endonucleases impart to their host phage remains unclear. Recently, a homing endonuclease was shown to facilitate interference competition between coinfecting phages, exemplifying a possible selective advantage to these proteins(*19*).

**Fig. 3.**
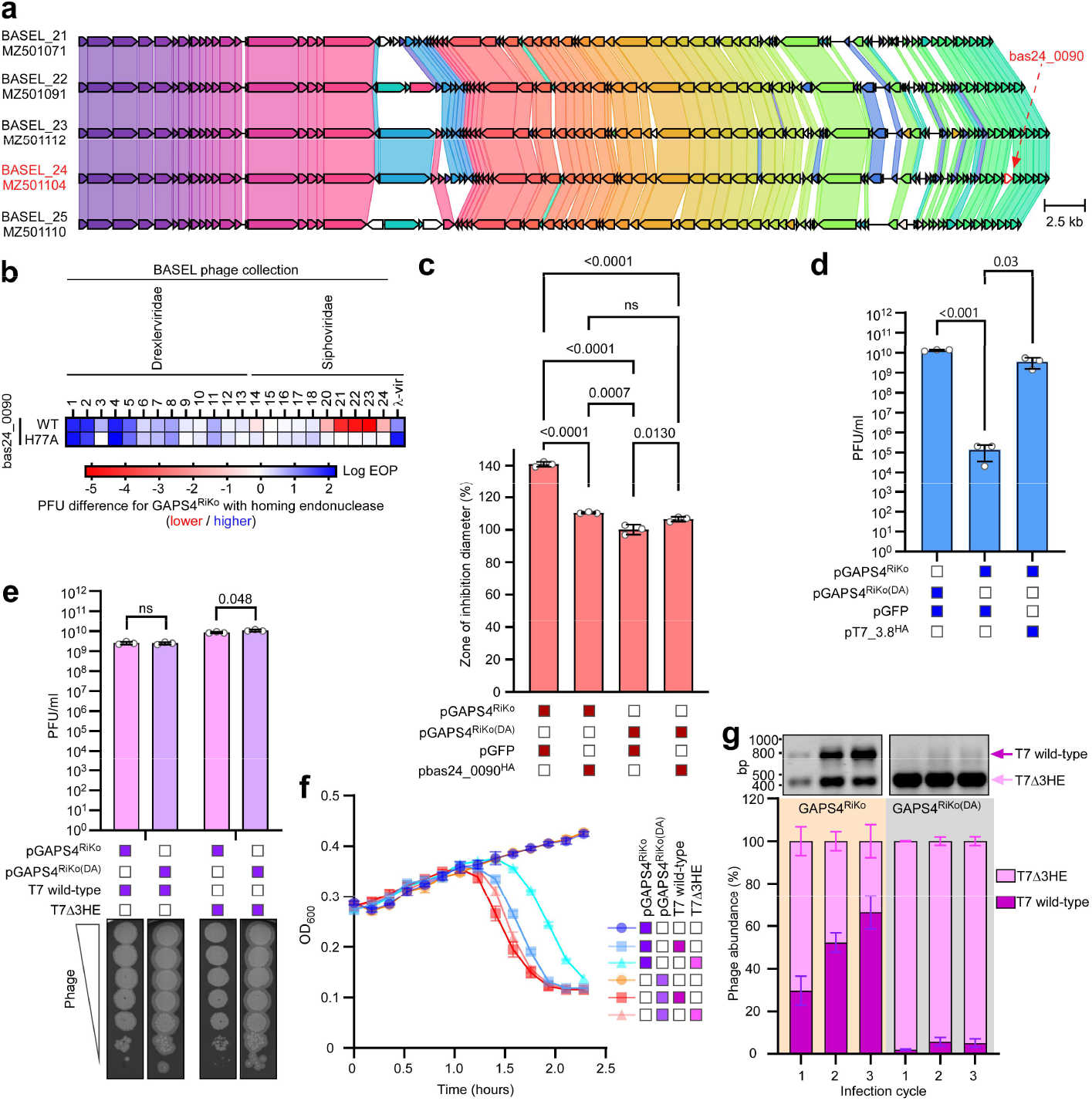
Homing endonucleases abrogate GAPS4-mediated defense. **(a)** Comparison of BASEL_21 – BASEL_25 phage genomes. Colors denote homology between the encoded protein sequences. The strain name and RefSeq accession are denoted. The BASEL_24-specific *bas24_0090* is denoted by a red arrow. Genome similarity was visualized using Clinker. **(b)** The efficiency of plating (EOP; shown as the order of magnitude difference in plaque numbers) was determined for *E. coli* harboring a low-copy plasmid encoding GAPS4^RiKo^ and a plasmid for the arabinose-inducible expression of the indicated *bas24_0090* form, when challenged with the indicated coliphages, compared to *E. coli* containing GAPS4^RiKo^ and a plasmid expressing GFP. The data shown are the average of 3 independent experiments. **(c)** Sensitivity of *E. coli* strains harboring a plasmid expressing GAPS4^RiKo^ (pGAPS4^RiKo^) or its catalytic mutant (pGAPS4^RiKo(DA)^) together with a plasmid for the arabinose-inducible expression of GFP or bas24_0090^H77A^ to nalidixic acid, as measured by the zone of inhibition diameter around a filter disc containing the antibiotics. Statistical significance between samples was calculated using One-way ANOVA with Tukey multiple comparisons correction. Data are shown as mean ± SD of 3 independent experiments. **(d)** Plaque-forming units (Pfus) of λ-vir phage upon infection of *E. coli* strains containing plasmids for the expression of the indicated GAPS4^RiKo^ from its native promoter and the arabinose-inducible expression of GFP or T7_3.8^H34A^. Statistical significance between samples was calculated using One-way ANOVA with Dunnett multiple comparisons correction. Data are shown as mean ± SD of 3 independent experiments. **(e)** Pfus of T7 phages upon infection of *E. coli* strains containing plasmids for the expression of the indicated GAPS4^RiKo^ from its native promoter. Statistical significance between samples was calculated using an unpaired, two-tailed Student *t*-test. **(f)** Growth of *E. coli* strains (measured as optical density at 600 nm; OD_600_) containing plasmids for the expression of the indicated GAPS4^RiKo^ form and infected or not with the indicated T7 phage at a multiplicity of infection (MOI) of 0.001. Data are shown as mean ± SD of 3 independent experiments. **(g)** Competition between wild-type T7 and T7Δ3HE phages that were initially added at a 1:10 ratio to equivalent populations of *E. coli* strains harboring a plasmid expressing GAPS4^RiKo^ or its catalytic mutant (GAPS4^RiKo(DA)^). The percentage of the phage population at the end of consecutive infection cycles was determined by PCR amplification. A representative gel is shown on top. Data are shown as mean ± SD of 3 independent experiments. In c and e: ns, no significant difference (*P* > 0.05).

To determine whether bas24_0090 hinders the protection provided by GAPS4 against invading phages, we expressed it from an arabinose-inducible plasmid in *E. coli* cells. When comparing the EOP of various lytic phages upon infection of GAPS4^RiKo^-containing *E. coli* strains harboring a plasmid expressing bas24_0090 or an empty plasmid, we observed a dramatic effect of bas24_0090; most of the examined phages regained their ability to infect *E. coli* (i.e., form plaques) in the presence of GAPS4 (Fig. 3b), indicating that this homing endonuclease hinders GAPS4-mediated protection. Notably, bas24_0090 appears to have the opposite effect on BASEL_21 – BASEL_25 phages, further hindering their ability to infect *E. coli*. This is to be expected, since these phages likely harbor the specific cleavage site of bas24_0090 in their genomes, thus leading to a phenotype resembling the recently reported homing endonuclease-mediated interference competition(*19*).

Next, we set out to identify the mechanism by which bas24_0090 abrogates GAPS4-mediated defense. First, we asked whether nuclease activity is required. A mutation in the predicted HNH active site histidine 77 to alanine (H77A) retained the ability to hinder GAPS4-mediated protection against phage attacks at levels comparable to those of its wild-type counterpart (Fig. 3b). However, it was unable to hinder BASEL_21 – BASEL_25 infection, confirming that this is a catalytically inactive mutant unable to mediate interference competition. Importantly, the expression of the catalytic mutant form of bas24_0090 did not affect the transcript level of GAPS4^RiKo^, confirming that it does not abrogate GAPS4-mediated defense simply by downregulating GAPS4 expression (Fig. S7).

The above results indicated that GAPS4 cleaves DNA, even in the absence of a phage, if it senses DNA breaks (e.g., in the presence of nalidixic acid (Fig. 2c-d)). We reasoned that this ability can be used to determine whether the attenuating effect of the homing endonuclease stems from its activity on the incoming phage or on the bacterial host (e.g., inhibiting GAPS4 activity or masking DNA breaks). Remarkably, the expression of bas24_0090^H77A^ abolished the toxic effect of GAPS4^RiKo^ in the presence of nalidixic acid (Fig. 3c). This result reveals that the homing endonuclease abrogates GAPS4-mediated toxicity regardless of the attacking phage, possibly by preventing GAPS4 from sensing or cleaving the damaged bacterial DNA.

### Homing endonucleases confer a competitive advantage to their host phage

Infection by the T7 phage is only mildly affected by the presence of GAPS4 (Fig. 1d). We hypothesized that T7 homing endonucleases play a role in hindering GAPS4-mediated defense. To investigate this possibility, we examined the effect of another HNH-containing homing endonuclease encoded by phage T7, T7_3.8 (NP_041974.1; 46% identity over 17% of the sequence compared to bas24_0090), on lytic phage λ attacks. Remarkably, the ectopic expression of a catalytically inactive T7_3.8^H34A^ from a plasmid dramatically hindered GAPS4-mediated protection, as evidenced by the increase in plaque numbers compared to a plasmid for GFP expression (Fig. 3d). These results demonstrate that diverse homing endonucleases can hinder GAPS4-mediated protection.

Surprisingly, infection of *E. coli* with the wild-type T7 phage or a mutated phage in which we deleted three homing endonucleases, including T7_3.8 (T7Δ3HE), resulted in a comparable number of plaques whether GAPS4^RiKo^ or its inactive mutant were expressed (Fig. 3e). We attribute this to additional yet unknown protection mechanisms possibly harbored by the T7 phage, which mask the deletion of the homing endonucleases in the phage context. Nevertheless, we did observe a difference in the plaque size between the wild-type and mutant phages, as the latter produced smaller plaques in the presence of the wild-type GAPS4^RiKo^ compared to the GAPS4^RiKo(DA)^ inactive mutant (Fig. 3e), suggesting that the homing endonucleases are required for normal phage growth in the presence of GAPS4. Indeed, we observed a delay in the decline of the bacterial population upon phage infection when *E. coli* harboring GAPS4^RiKo^ were challenged with T7Δ3HE (Fig. 3f); notably, only a mild difference between the effect of the wild-type and mutant T7 phages was observed when *E. coli* harboring the inactive GAPS4^RiKo(DA)^ were challenged, confirming that the phage titers were comparable and that the delay was mainly GAPS4-dependent (Fig. 3f). Strikingly, when we challenged *E. coli* with a mixture of the wild-type T7 and its mutant T7Δ3HE, the wild-type phage quickly outcompeted the mutant in the presence of GAPS4^RiKo^ but not in the presence of its inactive GAPS4^RiKo(DA)^ (Fig. 3h). Collectively, these results demonstrate a previously unknown role for phage-borne homing endonucleases in attenuating bacterial immunity, thus enhancing the competitive fitness of their host phage.

## Discussion

Numerous anti-phage defense systems have been described in recent years(*5*), yet in most cases, their trigger and activity remain unknown, as are potential counter-acting phage anti-defense determinants(*2*). Here, we describe GAPS4, a widespread anti-phage defense system that cleaves DNA upon sensing DNA breaks. We also uncover another layer of the intricate relationship between bacteria and their rivals by proposing a new role for phage-encoded homing endonucleases. Although mainly considered as selfish elements until recently, we show that homing endonucleases enhance phage fitness by attenuating GAPS4-mediated immunity.

GAPS4 is a DNase that is activated upon sensing DNA breaks induced by attacking phages, antibiotic compounds, or antibacterial toxins. It degrades both the host and the phage DNA, indicating that, unlike several other anti-phage defense systems(*20*), GAPS4 does not require DNA modification that distinguishes between self and non-self DNA to recognize its target. Therefore, GAPS4 is an abortive defense system. Sensing DNA breaks or linear DNA to activate the nuclease activity of a PD-(D/E)xK effector domain appears to be a common theme in anti-phage defense systems. For example, Shedu(*21, 22*) and Hachiman(*23*) were shown to target DNA upon sensing dsDNA ends or aberrant DNA states to nick or cleave dsDNA, respectively, although the sensor and activation mechanisms differ from those of GAPS4.

The proposed activation mechanism for GAPS4 is in agreement with the recent observation that GAPS4 incurs an expression-controlled trade-off between the benefit of protection range and the cost of autoimmunity(*24*). When overexpressed, it is more likely to sense random DNA breaks in the bacterial genome, thereby degrading the genomic DNA, leading to cell death. Furthermore, GAPS4 and possibly other anti-phage defense systems that sense DNA damage are potential double-edged swords. As we have demonstrated, they can sensitize bacteria to antagonistic attacks that induce DNA breaks. Another anti-phage defense system, CBASS, was recently shown to increase sensitivity to antifolate antibiotics(*25*). Although they employ different mechanisms of action, the phenomenon of anti-phage defenses incurring a fitness trade-off when bacteria are faced with non-phage antagonism appears to be more common than anticipated(*26*).

Homing endonucleases have long been considered selfish mobile elements found in many phage genomes(*17*), yet their contribution to phage fitness was poorly understood. Recently, Birkholz et al. reported that a homing endonuclease provides a relative fitness advantage during phage co-infection, by mediating inter-phage antagonism termed interference competition(*19*). Here, we report a new role for phage-encoded homing endonucleases in attenuating bacterial immunity to enhance phage fitness. We show that two homing endonucleases from different phages hinder GAPS4-mediated protection when ectopically over-expressed in the target bacterium. In contrast, these homing endonucleases appear to have a mild effect on GAPS4-mediated defense when expressed naturally from a phage genome during infection. Rather than abolishing GAPS4 protection (i.e., increasing the EOP), they attenuate it (i.e., accelerate the infection cycle). We assume that additional phage-encoded factors confer protection against GAPS4, either through specific or non-specific mechanisms, which likely explains why the observed effect of the homing endonucleases is mild in the context of an infecting phage.

We show that a homing endonuclease also prevents GAPS4-mediated toxicity induced by nalidixic acid. Therefore, we conclude that rather than targeting or manipulating the incoming phage, homing endonucleases interfere with GAPS4’s ability to sense DNA breaks. Remarkably, their ability is independent of the predicted nuclease activity, suggesting that it relies on the DNA-binding properties of the homing endonucleases. Since two different homing endonucleases attenuated GAPS4-mediated protection, it is unlikely that they directly inhibit GAPS4 activity. Therefore, we hypothesize that homing endonucleases bind to regions containing DNA breaks and mask them from GAPS4. This could delay GAPS4 activation and thus cell suicide, allowing the incoming phage more time to complete the infection cycle and spread further in the bacterial population.

## Materials and Methods

### Strains and Media

For a complete list of strains used in this study, see Table S1. *Escherichia coli* strains were grown in lysogeny broth (LB; 1% [w/v] tryptone, 0.5% [w/v] yeast extract, and 0.5% [w/v] NaCl) or 2xYT (1.6% [w/v] tryptone, 1% [w/v] yeast extract, and 0.5% [w/v] NaCl) at 37°C. *Vibrio parahaemolyticus* strains were grown in marine lysogeny broth (MLB; LB containing 3% [w/v] NaCl) and on marine minimal media (MMM) agar plates (1.5% [w/v] agar, 2% [w/v] NaCl, 0.4% [w/v] galactose, 5 mM MgSO_4_, 7 mM K_2_SO_4_, 77 mM K_2_HPO_4_, 35 mM KH_2_PO_4_, and 2 mM NH_4_Cl) at 30°C. Media were supplemented with 1.5% (w/v) agar to prepare solid plates. When required, media were supplemented with 35 or 10 µg ml^−1^ chloramphenicol (for *E. coli* and *V. parahaemolyticus*, respectively), 50 or 250 µg ml^−1^ kanamycin (for *E. coli* and *V. parahaemolyticus*, respectively), or 100 µg ml^−1^ ampicillin to maintain plasmids. To induce the expression from P*bad* promoters, 0.025%, 0.04%, or 0.2% (w/v) L-arabinose was added to the media, as indicated.

### Plasmid construction

For a complete list of plasmids and primers used in this study, see Table S2 and Table S3, respectively. Plasmids were constructed with standard molecular biology techniques using the Gibson Assembly method(*27*), unless otherwise mentioned. The Gibson Assembly master mix was obtained from NEB (E2611S). DNA fragments were amplified by PCR from bacterial genomic DNA or from bacteriophage genomic DNA, and Gibson Assembly ligations were carried out according to the manufacturer’s instructions.

### Construction of T7 mutant phages

T7Δ3HE was constructed using homologous recombination, as previously described(*28*). Briefly, Neb5α bacteria harboring pGEM^2.8OH^-*trxA*, a plasmid encoding *trxA-FLP* flanked by sequences homologous to those upstream and downstream of T7 gene 2.8, were grown overnight in LB supplemented with ampicillin at 37°C. The cells were diluted 1:100 in 3 ml of fresh LB supplemented with ampicillin and grown at 37°C. Upon reaching OD_600_ of ∼0.5, the cells were infected with wild-type T7 phage. The lysate, harboring some recombinant phages that replaced gene 2.8 with *trxA*, was mixed with *E. coli* BW25113 Δ*trxA* strain lacking *trxA* in LB containing 0.7% (w/v) molten agar, and poured onto a 1.5% (w/v) agar plate to select the recombinant phage T7Δ2.8:: trxA-FLP. Plaques were then streak-purified on BW25113 Δ*trxA*. The isolated phage T7Δ2.8:: *trxA-FLP* was used to infect *E. coli* BW25113 cells harboring pAC-ff-*FLP* encoding FLP recombinase to flip out the *trxA*. The lysates, harboring some recombinant phages where the *trxA* was flipped out, were mixed with *E. coli* BW25113. Several plaques were picked and spotted on 1.5% (w/v) agar plates containing a lawn of either *E. coli* BW25113 or *E. coli* BW25113 Δ*trxA* cells. To isolate T7Δ2.8 phage with flipped out *trxA*, we relied on the fact that these phages should form plaques on *E. coli* BW25113 but not on *E. coli* BW25113 Δ*trxA*. Then, T7Δ2.8 phage was used to infect Neb5α harboring pGEM^7.7OH^-*trxA*, a plasmid encoding *trxA-FLP* flanked by sequences homologous to those upstream and downstream of T7 gene 7.7. T7Δ2.8Δ7.7 phages were isolated using the methodology described above. Finally, to engineer T7Δ3HE phage (T7Δ2.8Δ7.7Δ3.8:: *trxA-FLP*), T7Δ2.8Δ7.7 was used to infect Neb5α harboring pGEM^3.8OH^-*trxA*, a plasmid encoding *trxA-FLP* flanked by sequences homologous to those upstream and downstream of T7 gene 3.8. T7Δ3HE phage was then isolated as described above.

### Plaque assays

The phages used in this study are listed in Table S4. Phages were propagated on *E. coli* K12 MG1655 ΔRM(*29*). To determine the effect of GAPS4^RiKo^ against coliphages T4, T5, T7, λ-vir, P1-vir, and 68 phages belonging to the BASEL collection(*29*), *E. coli* K12 MG1655 ΔRM strains harboring the pGAPS4^RiKo^ plasmid or an empty pC-0_v5 plasmid were grown overnight in LB supplemented with ampicillin at 37°C. Then, 200 μl of each bacterial culture were mixed with 7 ml of 0.7% (w/v) molten agar supplemented with 10 mM MgSO_4_ and 5 mM CaCl_2_. The mixtures were poured onto 1.5% (w/v) agar plates supplemented with ampicillin, 10 mM MgSO_4_, and 5 mM CaCl_2_, and the plates were left to dry. Tenfold serial dilutions of all the phages were prepared, and 7.5 μl of each dilution were spotted onto the dried plates. The plates were incubated overnight at 37°C. The following day, the plaques were counted and the plaque-forming units (p.f.u.s ml^−1^) were calculated. For dilution spots in which no individual plaques were visible but a faint zone of lysis was observed, the dilution was considered as having ten plaques, as previously described(*30*). A similar plaque assay was performed with *E. coli* MG1655 strains containing either pGAPS4^RiKo^ or pGAPS4^RiKo(D49A)^ using λ-vir, T7, or T7Δ3HE. Plaque assays with λ-vir phage infecting *E. coli* MG1655 strains containing pBAD33.1-based plasmids encoding the indicated GAPS4^MAVP-R^ versions were performed on LB agar supplemented with chloramphenicol, 0.1% (w/v) L-arabinose (to induce the expression from the P*bad* promoter), and 10 mM MgSO_4_ at 37°C. This protocol was also used to investigate the ability of the indicated phages to form plaques on *E. coli* K12 MG1655 ΔRM strains containing pGAPS4^RiKo^ together with a plasmid for expression of a homing endonuclease or a fluorescent protein control, except the media were supplemented with ampicillin, kanamycin, 0.04% (w/v) L-arabinose, 10 mM MgSO_4_, and 5 mM CaCl_2_.

### Lysogenization and viability assays

*E. coli* MG1655 strains harboring pGAPS4^RiKo^ or pGAPS4^RiKo(D49A)^ were grown overnight in LB supplemented with ampicillin at 37°C. Overnight cultures were diluted 1:100 in 10 ml of fresh LB supplemented with ampicillin, 10 mM MgSO_4_, and 0.2% (w/v) D-maltose, and grown to an OD_600_ of ∼0.5. The cells were then infected with λ phage CI-857 containing a kanamycin-resistance cassette(*31*) at a multiplicity of infection (MOI) of ∼0.1 or ∼5, and incubated for 120 minutes at 30°C. Tenfold serial dilutions of these infected cultures were spotted onto 1.5% (w/v) agar plates supplemented with kanamycin and ampicillin (to select for lysogens) or only ampicillin (to determine the total number of viable cells), and the plates were incubated overnight at 30°C. The colonies were then counted, and the CFU ml^−1^ was calculated.

### Transformation into λ lysogen assays

*E. coli* BW25113 cells lysogenized with λ phage CI857 containing a kanamycin-resistance cassette were grown overnight in LB supplemented with kanamycin at 30°C. The cells were diluted 1:100 in 10 ml of fresh LB supplemented with kanamycin and grown at 30°C. Upon reaching an OD_600_ of ∼0.5, the cells were harvested and washed twice with chilled double-distilled H_2_O (DDW) and resuspended in 150 μl of chilled DDW. The cells were then electroporated with equal molar amounts of pGAPS4^RiKo^ or pGAPS4^RiKo(D49A)^ and then diluted and spotted onto 1.5% (w/v) agar plate supplemented with kanamycin and ampicillin. The plates were incubated overnight at 30°C. The next day, the transformed colonies were counted, and the CFU ml^−1^ was calculated.

### Transduction assays

Transduction assays were carried out as previously described(*32*). Briefly, *E. coli* BW25113 cells harboring the T7-packaging signal-containing pIYPE69 plasmid were grown overnight in LB supplemented with kanamycin at 37°C. The cells were diluted 1:100 in fresh LB supplemented with kanamycin and grown at 37°C. Upon reaching an OD_600_ of ∼0.5, the cells were infected with wild-type T7 phage at an MOI of 2. Infected cells were grown until lysis occurred (1.5-2 hours). Then, 200 µl of chloroform were added, and the lysates were centrifuged to clear cell debris. The resulting supernatants comprised a mixture of T7 phage particles with either a T7 genome or a pIYPE69 plasmid containing a T7-packaging signal. For the transduction assay, *E. coli* BW25113 Δ*trxA* strains harboring pGAPS4^RiKo^ or pGAPS4^RiKo(D49A)^ were grown overnight in LB supplemented with ampicillin at 37°C. Overnight cultures were diluted 1:100 in 3 ml of fresh LB supplemented with ampicillin and grown to an OD_600_ of ∼0.5. The cells were then mixed at a 1:1 ratio (v/v) with tenfold serial dilutions of the prepared phage lysates described above. Transduction was carried out for 1 hour at 37°C with shaking. Then, the cells were spotted onto 1.5% (w/v) agar plate supplemented with kanamycin and ampicillin, and incubated overnight at 37°C. The next day, the transduced colonies were counted, and the CFU ml^−1^ was calculated.

### Quantitative real-time PCR assays

To monitor genomic DNA stability, *E. coli* MG1655 cells harboring pGAPS4^RiKo^ or pGAPS4^RiKo(D49A)^ were grown overnight in LB supplemented with ampicillin at 37°C. Overnight cultures were diluted 1:100 in 10 ml of fresh LB supplemented with ampicillin, 10 mM MgSO_4_, and 0.2% (w/v) D-maltose, and grown to an OD_600_ of ∼0.5. The cells were then infected with λ phage CI-857 containing a kanamycin-resistance cassette at an MOI of ∼1 and incubated for 90 minutes at 30°C. Cells equivalent to 0.5 OD_600_ units were harvested and washed twice with LB. The genomic DNA was extracted using the Presto^TM^ Mini gDNA isolation kit (Geneaid) and diluted 1:10 in DDW. For real-time (RT)-PCR, an equal amount of gDNA was mixed (in triplicate) with forward and reverse primers (400 nM each; detailed below) and with 1x AzuraView GreenFast qPCR Blue Mix. The RT-PCR and subsequent analyses were carried out using a QuantStudio 3 real-time PCR system and software (Applied Biosystems). The primer pair TM623F and TM634R was used for the detection of λ phage genomic DNA, and the primer pair TM110F and TM110R was used for the detection of bacterial genomic DNA (Table S3).

To monitor GAPS4 transcription, *E. coli* MG1655 cells harboring pbas24_0090^H77A^ or pGFP, together with either pGAPS4^RiKo^ or pGAPS4^RiKo(D49A)^, were grown overnight in LB supplemented with ampicillin and kanamycin at 37°C. Overnight cultures were diluted 1:100 in 10 ml of fresh LB supplemented with ampicillin and kanamycin, and grown up to an OD_600_ of ∼0.5. L-arabinose (0.2% [w/v]) was added to the cultures to induce expression from P*bad* promoters, and cells were grown for 4 additional hours at 37°C. Cells equivalent to 1 OD_600_ units were harvested, and RNA was isolated using a Bacterial RNA kit (Biomiga), following the manufacturer’s instructions. Complementary DNA (cDNA) was synthesized from isolated RNA (1 µg) using the AzuraQuant II cDNA synthesis kit (Azura genomics), following the manufacturer’s instructions. For RT-PCR, 2 ng template cDNA were mixed (in triplicate) with forward and reverse primers (400 nM each; detailed below) and with 1x AzuraView GreenFast qPCR Blue Mix. The RT-PCR and subsequent analyses were carried out using a QuantStudio 3 real-time PCR system and software (Applied Biosystems). 16s rRNA was used as the endogenous control, and the differential gene expression, reported as the fold change, was analyzed as the ΔCt values. The primer pair TM769F and TM769R was used to monitor GAPS4^RiKo^, and the primer pair TM110F and TM110R was used to monitor 16s rRNA (Table S3).

### In vivo DNA stability assays

To determine the effect of phage infection on DNA stability, *E. coli* MG1655 strains harboring pGAPS4^MAVP-R^or pGAPS4^MAVP-R(D59A)^were grown overnight in LB supplemented with chloramphenicol and 0.2% (w/v) D-glucose at 37°C. Overnight cultures were washed twice, diluted 1:100 in 10 ml of fresh LB supplemented with chloramphenicol, and grown for an hour at 37°C. L-arabinose (0.04% [w/v]) was added to the cultures to induce expression from the P*bad* promoter, and cells were grown for an additional hour at 37°C. The cells were then infected with λ-vir or Basel_01 phage at an MOI of ∼10, and the cultures were incubated for 90 minutes at 37°C with shaking. Then, cells equivalent to 1 OD_600_ units were collected from each sample, and genomic DNA was isolated using the Presto^TM^ Mini gDNA isolation kit. Equal volumes of genomic DNA were then resolved on a 0.8% (w/v) agarose gel.

To determine the effect of nalidixic acid on bacterial genomic DNA in the presence of GAPS4^MAVP-R^, *E. coli* MG1655 strains harboring pGAPS4^MAVP-R^or pGAPS4^MAVP-R(D59A)^were grown overnight in LB supplemented with chloramphenicol and 0.2% (w/v) D-glucose at 37°C. Overnight cultures were washed twice, diluted 1:100 in 10 ml of fresh LB supplemented with chloramphenicol, and grown for 45 minutes at 37°C. L-arabinose (0.04% [w/v]) was added to the cultures to induce expression from the P*bad* promoter, and cells were grown for 45 additional minutes at 37°C. Next, nalidixic acid was added at a final concentration of 5 μg ml^−1^, and the cultures were incubated for 90 minutes at 37°C with shaking. Genomic DNA was extracted from cells equivalent to 1.0 OD_600_ units and resolved on a 0.8% (w/v) agarose gel. A similar experiment was carried out to study the effect of nalidixic acid on bacterial genomic DNA in the presence of GAPS4^RiKo^, with *E. coli* MG1655 cells harboring pGAPS4^RiKo^ or pGAPS4^RiKo(D49A)^. Overnight cultures were diluted 1:100 in 10 ml of fresh LB supplemented with ampicillin and grown for 90 minutes at 37°C. Nalidixic acid was added at a final concentration of 5 μg ml^−1^, and genomic DNA was extracted from cells equivalent to 1 OD_600_ units following incubation for 60, 120, and 180 minutes at 37°C with shaking, and resolved on a 0.8% (w/v) agarose gel.

### Microscopy

To visualize the effect of nalidixic acid on cellular DNA content in the presence of GAPS4^MAVP-R^ using fluorescence microscopy, *E. coli* MG1655 cells harbouring pGAPS4^MAVP-R^ or pGAPS4^MAVP-R(D59A)^ were grown overnight at 37°C in LB supplemented with chloramphenicol and 0.2% (w/v) D-glucose. Overnight cultures were washed twice and then diluted 1:100 in 3 ml of fresh LB supplemented with chloramphenicol and incubated at 37°C for 45 minutes. Cells were then induced with 0.025% arabinose and incubated for 45 additional minutes. Next, nalidixic acid was added at a final concentration of 5 μg ml^−1^, and the cells were incubated for 75 minutes. Following incubation, the cells were washed twice with M9 media and stained for 10 minutes at room temperature with Hoechst 33342 (Invitrogen) DNA dye at a final concentration of 1 μg μl^−1^. The cells were then washed twice and resuspended in 10 μl of M9 media. To visualize the cell wall of *E. coli*, cells were incubated with Wheat Germ Agglutinin Alexa Fluor 488 Conjugate (Biotium; Catalogue no. 29022–1) at a final concentration of 0.1 mg ml^−1^, and incubated for 10 minutes at room temperature. Next, 2 μl of each cell suspension were spotted onto 1% (w/v) LB-agarose pads, which were placed face-down in 35 mm glass-bottom Cellview cell culture dishes. The imaging was carried out using a Nikon Eclipse Ti2-E inverted motorized microscope equipped with a CFI PLAN apochromat DM 100X oil lambda PH-3 (NA, 1.45) objective lens, Lumencor SOLA SE II 395 light source, and DS-QI2 mono cooled digital microscope camera (16 MP). An ET-EGFP filter set (#49002) was used to visualize the Alexa Fluor 488 signal, and an ET-DAPI (#49028) was used to visualize the Hoechst signals. The captured images were further processed using Fiji ImageJ suite(*33*).

### Bacterial competition assays

Bacterial competition assays were performed as previously described(*34*), with minor modifications. Briefly, bacteria were grown overnight in appropriate media supplemented with antibiotics to maintain plasmids: *V. parahaemolyticus* attacker strains in MLB and *E. coli* IYB5101 prey in 2xYT. Overnight cultures were normalized to OD_600_ = 0.5 and mixed at a 1:1 ratio (attacker:prey). Triplicates of the mixtures were spotted onto MLB agar plates supplemented with 0.1% (w/v) L-arabinose, and incubated for 4 hours at 30°C. Samples from the 0-hour timepoint and samples recovered from the assay plates after 4 hours were plated on selective media to quantify the viability (colony-forming units; CFU) of the prey strains. The experiments were performed three times with similar results; results from a representative experiment are shown.

### Antibiotic sensitivity assays

Zone of inhibition assays were performed using the soft agar overlay technique. *E. coli* MG1655 strains harboring the indicated plasmids were grown overnight in 2xYT supplemented with antibiotics to maintain the plasmids. Overnight cultures were then diluted 1:100 into fresh media and grown to OD_600_ ∼0.9. Then, the cultures were mixed to a final concentration of 0.08 or 0.05 OD_600_ (for 130 mm square plates or 90 mm round plates, respectively) with 0.5% (w/v) molten agar lacking or supplemented with 0.025% (w/v) L-arabinose, which was added when inducible expression from a P*bad* promoter was required. The mixtures were poured onto 1.5% (w/v) LB agar plates, lacking or supplemented with 0.025% (w/v) L-arabinose, as described above, and the plates were left to dry for an hour at room temperature. Next, sterile Whatman paper discs (5 mm diameter) were placed on the surface. Each disc was spotted with 7 µl of antibiotic solution, delivering final amounts of 30 µg nalidixic acid, 10 µg gentamicin, 5 µg trimethoprim, 5 µg rifampicin, or 30 µg novobiocin. The plates were incubated at 37°C, and the diameter of each zone of inhibition around a disc was measured after 16 hours using the ImageJ software(*33*). Each assay included three biological replicates with three technical replicates per condition. The experiments were performed at least twice with similar results; results from a representative experiment are shown.

For spotting assays, *E. coli* MG1655 cells containing pGAPS4^MAVP-R^ or pGAPS4^MAVP-R(D59A)^ were grown overnight in LB supplemented with chloramphenicol and 0.2% (w/v) D-glucose at 37°C. Overnight cultures were washed twice, diluted 1:100 in 3 ml of fresh LB supplemented with chloramphenicol, and grown for 45 minutes at 37°C. L-arabinose (0.04% [w/v]) was added to the cultures to induce expression from the P*bad* promoter, and the cells were grown for 45 additional minutes at 37°C. Then, tenfold serial dilutions were spotted onto 1.5% (w/v) agar plate supplemented with chloramphenicol, L-arabinose (0.04% [w/v]), and the indicated concentrations of antibiotics. The plates were incubated overnight at 37°C.

### Phage T7 infection assays

To monitor bacterial growth upon infection with wild-type T7 or the T7Δ3HE phage, *E. coli* MG1655 cells harboring pGAPS4^RiKo^ or pGAPS4^RiKo(D49A)^ were grown overnight at 37°C in LB supplemented with ampicillin. Overnight cultures were diluted 1:100 in 10 ml of fresh LB and grown to an OD_600_ of ∼0.5. The phages were then added to the bacterial cultures at a multiplicity of infection (MOI) of 0.001, and 200 µl of cells were transferred into 96-well plates in triplicate. Cells were grown under continuous shaking (205 rpm) in a Tecan Infinite M Plex plate reader at 37°C. OD_600_ readings were acquired every 10 minutes.

For the phage competition assay, *E. coli* MG1655 cells harboring pGAPS4^RiKo^ or pGAPS4^RiKo(D49A)^ were grown overnight at 37°C in LB supplemented with ampicillin. Overnight cultures were diluted 1:100 in 10 ml of fresh LB and grown to an OD_600_ of ∼0.5. The competition cycle was initiated by infecting the logarithmic-phase bacterial culture with a 1:10 ratio of wild-type T7 and T7Δ3HE phage, respectively, at an MOI of 0.001. The infected cultures were grown at 37°C until complete lysis was observed. The obtained lysates were diluted 1:266 into freshly growing, naive logarithmic phase bacterial cultures for the next infection cycle. The relative abundance of wild-type T7 and T7Δ3HE in each cycle was determined by PCR amplification of the T7 genomic region flanking gp2.8 using primers 105F and SM24R8 (Table S3). The expected amplified products from wild-type T7 and T7Δ3HE were 706 bp and 400 bp, respectively. Band intensities per infection cycle were quantified using the ImageJ software, and the relative band intensity of each phage was calculated using the following formula: [(band corresponding to wild-type T7 or T7Δ3HE)/(sum of the band intensities)] × 100. The data were plotted using GraphPad Prism.

### Protein purification

For purification of the GAPS4^MAVP-R^ complex, pMBP-GAPS4^MAVP-R^ was transformed into *E. coli* BL21 (DE3) cells and grown in 2 l of 2xYT supplemented with 50 μg ml^−1^ kanamycin at 37°C until they reached an OD_600_ ∼0.5. Then, GAPS4 expression was induced by adding 0.5 mM IPTG, and the cells were grown at 16°C for 18 hours. The cells were then harvested and resuspended in lysis buffer (20 mM HEPES–NaOH pH 7.5, 300 mM NaCl, 10 mM imidazole, 1 mM dithiothreitol [DTT], 1 mM phenylmethylsulfonyl fluoride [PMSF], 10% [v/v] glycerol). Sonication was used to lyse the cells, and the supernatant was collected after centrifugation. The cleared supernatant was loaded onto a 1 ml His-Trap column (GE Healthcare) and washed with 25 column volumes of wash buffer (20 mM HEPES– NaOH pH 7.5, 300 mM NaCl, 10 mM imidazole, 1 mM DTT, 1 mM PMSF, 5% [v/v] glycerol). The bound proteins were then eluted with elution buffer (20 mM HEPES– NaOH pH 7.5, 300 mM NaCl, 300 mM imidazole, 1 mM DTT, 5% [v/v] glycerol) and dialyzed overnight at 4°C against dialysis buffer (20 mM HEPES–NaOH pH 7.5, 300 mM NaCl, 10 mM imidazole, 1 mM DTT, 5% [v/v] glycerol) in the presence of TEV protease to remove the His-Maltose binding protein (MBP) tag. The cleaved proteins were again passed through a 1 ml His-Trap column, and the flow-through containing the cleaved proteins was collected. The proteins were concentrated using Amicon Ultra-4 Centrifugal Filter Units (10 kDa columns) and further purified in a Superdex 200 Increase 10/300 size-exclusion chromatography column (GE Healthcare) equilibrated with protein buffer (20 mM HEPES–NaOH pH 7.5, 150 mM NaCl, 1 mM DTT). The fractions containing the protein were pooled and concentrated using Amicon Ultra-4 Centrifugal Filter Units (10 kDa columns), snap frozen, and stored at −80°C.

### SEC-MALS analyses

Experiments were carried out using an analytical size-exclusion chromatography (SEC) column (Superdex 200 10/300 GL), pre-equilibrated with SEC buffer (20 mM HEPES–NaOH pH 7.5, 150 mM NaCl, 1 mM DTT). Purified GAPS4^MAVP-R^ complex (1-2 mg ml^−1^; 50 µl) was injected onto an HPLC system connected in-line to an 8-angle multi-angle light-scattering (MALS) detector and a differential refractive index (RI) detector (Wyatt Technology, CA, USA). RI and MALS data were analyzed using ASTRA software (Wyatt Technology) to determine the absolute molecular mass of the complex.

### AlphaFold structure predictions

GAPS4 complex structures were predicted using AlphaFold 3(*35*) (https://alphafoldserver.com/). The best model was selected and visualized using ChimeraX(*36*) version 1.7.1 together with its Predicted Aligned Error (PAE) plot.

### Comparative genomics analyses

Genbank files of the indicated phage genomes were downloaded from NCBI and aligned using the CLINKER(*37*) server (cagecat.bioinformatics.nl/tools/clinker) with a minimal alignment sequence identity of 0.3.

### Identification of GAPS4 homologs

Each of the *Vibrio* GAPS4a and GAPS4b (WP_055466293.1 and WP_055466294.1) sequences was used to build a profile hidden Markov model (pHMM) using hmmbuild from the HMMer suite(*38*) (version 3.4, Aug 2023). These pHMMs were used to search all bacterial genomes from the NCBI GenBank Whole Genome Sequencing (WGS) database(*39*) (downloaded on March 14, 2020) to detect adjacent and fused GAPS4a and GAPS4b proteins. The concatenated and fused sequences were clustered using CD-HIT(*40*) with 0.9 global sequence identity. To further de-duplicate sequences using taxonomy, up to one sequence per order per cluster was selected, and the rest were discarded. The remaining sequences were aligned using MAFFT(*41*) (mafft-linsi), which was then used to construct a maximum-likelihood phylogenetic tree using RAxML(*42*) (version 8.2.12) with the PROTGAMMALG evolutionary model. The tree was visualized using the interactive Tree Of Life (iTOL)(*43*) web server.

### Illustration of conserved residues using WebLogo

The protein sequences of GAPS4a and GAPS4b homologs were aligned using CLUSTAL Omega(*44*) in MEGA11(*45*). Aligned columns not found in representative proteins (WP_055466293.1 and WP_055466294.1) were discarded. The conserved residues were illustrated using the WebLogo server (weblogo.berkeley.edu)(*46*).

## Supporting information

Supplemental Figures, Tables, Files, Datasets, References

## Data Availability

All data related to the manuscript are available in the main text and the supplementary material files.

## Acknowledgments

We thank members of the Salomon and Qimron laboratories for helpful discussions and suggestions; Prof. Ulrich Dobrindt (University of Münster) for gifting us the *E. coli* RiKo 2299/09 strain; and Dr. Alexander Harms (ETH Zurich) for generously sharing the BASEL phage collection. D.S. received funding from the Israel Science Foundation (ISF grant number 1362/21) and the European Research Council – Horizon Europe research and innovation programme (grant number 101169966). U.Q. was supported by the European Research Council – Horizon 2020 research and innovation programme (grant number 818878). Y.H. was supported by the Israel Science Foundation (ISF grant number 1653/21). G.S. was funded in part by a fellowship from the Edmond J. Safra Center for Bioinformatics at Tel-Aviv University.

## Author Contributions

T.M., U.Q., and D.S. conceived the work; T.M., K.K., U.Q., and D.S. designed experiments; T.M. and K.K. carried out most of the experiments; R.M.R. and A.H. assisted in performing phage plaque assays; G.S. and D.B. performed the GAPS4 homology analyses; R.Y. and Y.H. performed the SEC-MALS analyses; U.Q. and D.S. acquired the funding and supervised the work; D.S. wrote the manuscript draft, and all authors edited and approved the paper.

