## Supplemental Figures, Tables, Files, Datasets, References for "Phage-encoded homing endonucleases attenuate bacterial immunity"

**Figures S1-S7**
**Tables S1-S4**
**Dataset S1**
**Files S1-S3**
**References**

### Supplementary Figures

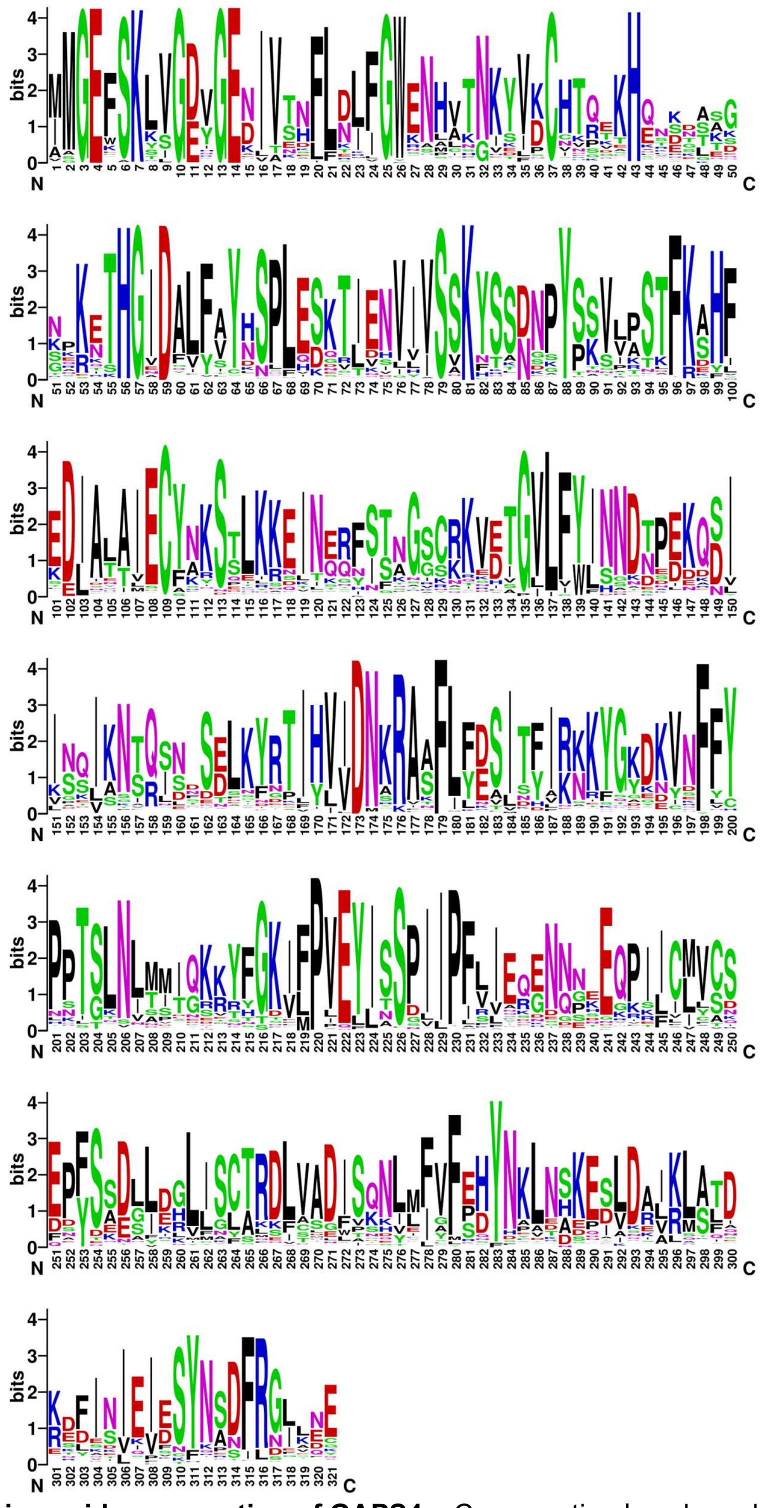

**Fig. S1. Amino acid conservation of GAPS4a.** Conservation logo based on a multiple sequence alignment of GAPS4a homologs. The position numbers correspond to the amino acids in WP_055466293.1.

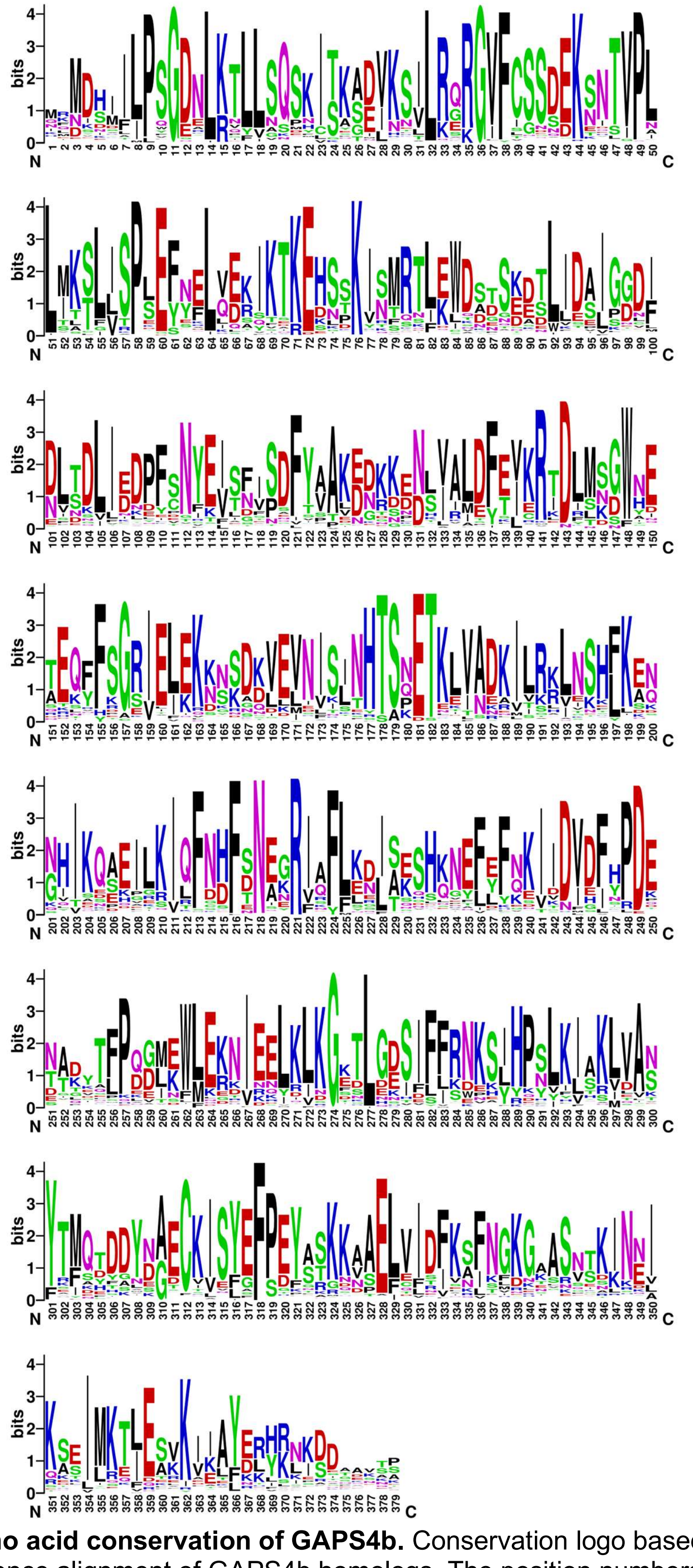

**Fig. S2. Amino acid conservation of GAPS4b.** Conservation logo based on a multiple sequence alignment of GAPS4b homologs. The position numbers correspond to the amino acids in WP_055466294.1.

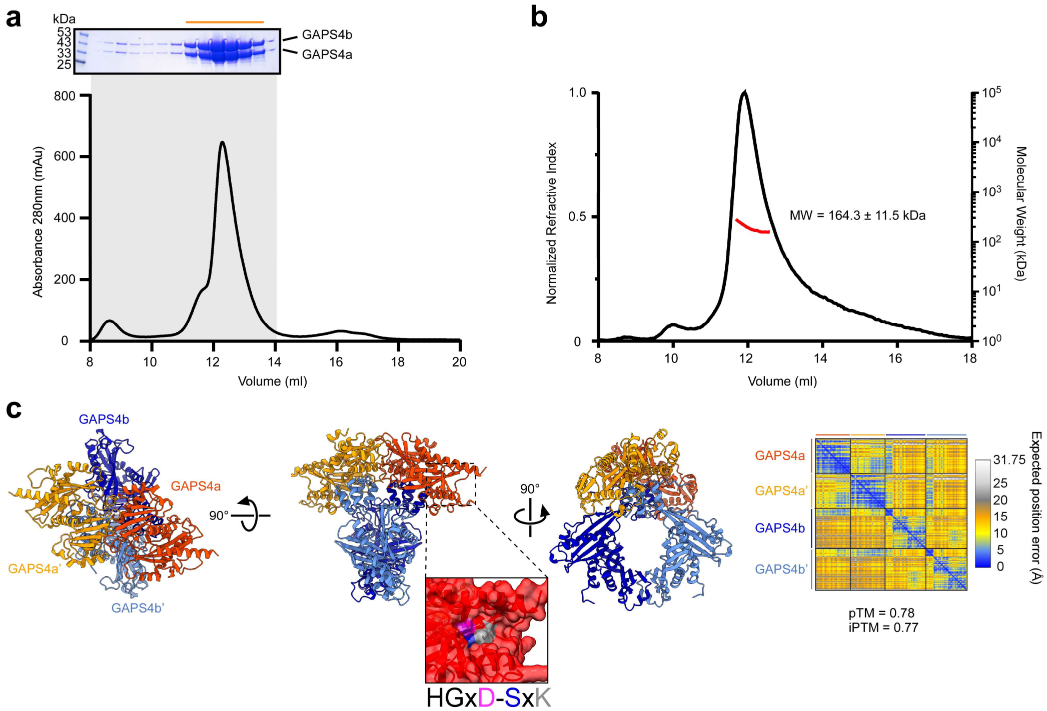

**Fig. S3. GAPS4$^{MAVP-R}$ is a heterotetramer complex. (a)** Size-Exclusion Chromatography (SEC) purification of GAPS4. Coomassie blue-stained SDS-PAGE of the eluted fractions denoted in gray is shown above. The fractions denoted by an orange line were pooled and used for subsequent analyses. **(b)** SEC-MALS analysis of purified GAPS4. The black line represents the normalized refractive index, and the red line represents the calculated molecular weight (MW) in kDa. **(c)** Structural model of GAPS4$^{MAVP-R}$ heterotetramer predicted by AlphaFold3. The AlphaFold Predicted Aligned Error (PAE) plot is shown on the right. The color key represents the expected position error for each pair of residues in Å units. Blue represents low predicted errors, indicating high confidence in the relative positions of those residues. The orange color indicates higher predicted errors, suggesting lower confidence. The rectangular inset is a close-up view of the predicted PD-(D/E)xK active site in GAPS4a.

**a**

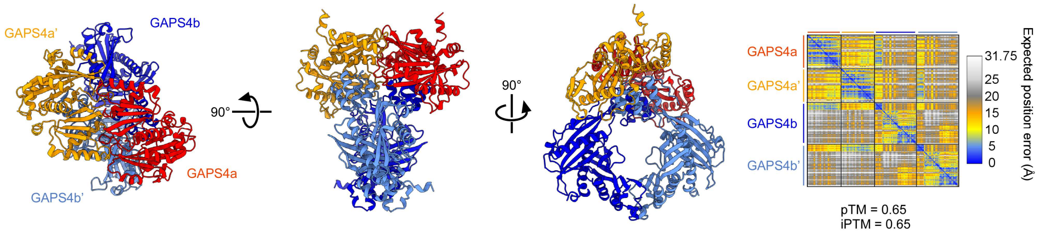

**b**

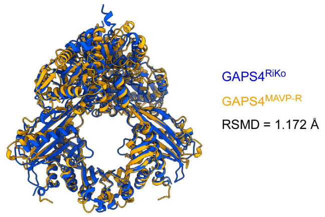

**Fig. S4. GAPS4$^{RiKo}$ is predicted to form a heterotetramer complex. (a)** Structural model of GAPS4$^{RiKo}$ predicted by AlphaFold3. The AlphaFold Predicted Aligned Error (PAE) plot is on the right. The color key represents the expected position error for each pair of residues in Å units. Blue represents low predicted errors, indicating high confidence in the relative positions of those residues. **(b)** Superimposition of the predicted GAPS4$^{RiKo}$ and GAPS4$^{MAVP-R}$ complexes.

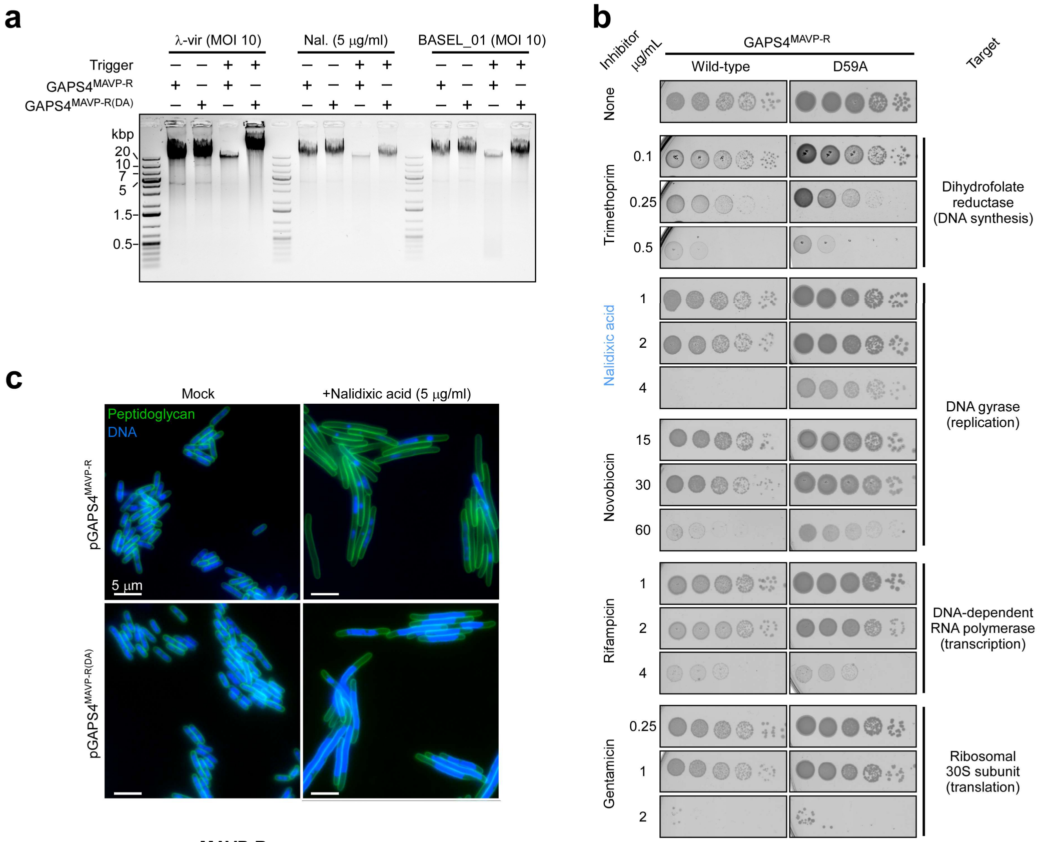

**Fig. S5. GAPS4$^{MAVP-R}$ is a DNase activated by phage infection and nalidixic acid.** **(a)** Genomic DNA stability after the addition of the indicated triggers to the media of *E. coli* cultures expressing the indicated GAPS4$^{MAVP-R}$ form from an arabinose-inducible plasmid (λ-vir phage, nalidixic acid [Nal.], or BASEL_01 phage). MOI, multiplicity of infection. The experiment was repeated three times with similar results. Results from a representative experiment are shown. **(b)** Growth of *E. coli* MG1655 strains expressing the indicated GAPS4$^{MAVP-R}$ forms from an arabinose-inducible plasmid on agar plates containing 0.04% (w/v) L-arabinose. The media were supplemented with the indicated inhibitors at the indicated final concentrations. **(c)** Sample fluorescence microscopy images of *E. coli* strains expressing the indicated GAPS4$^{MAVP-R}$ form from an arabinose-inducible plasmid after 75 minutes of incubation with 5 µg/ml nalidixic acid. Peptidoglycan (green) and DNA (blue) are visualized. Scale bar = 5 µm. The experiment was repeated three times with similar results.

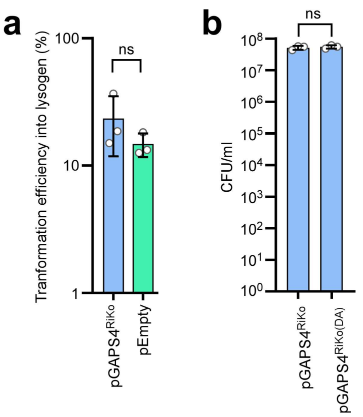

**Fig. S6. λ phage lysogens and T7 phage-mediated transduction do not induce GAPS4-mediated cell suicide. (a)** Transformation efficiency (percentage) of an empty plasmid (pEmpty) and a plasmid expressing GAPS4$^{RiKo}$ (pGAPS4$^{RiKo}$) into a λ lysogen *E. coli* strain. **(b)** Colony-forming units (CFU) of *E. coli* BW25113 Δ*trxA* containing a plasmid expressing the indicated GAPS4$^{RiKo}$ form, transduced with a T7 phage-based particle containing a plasmid conferring kanamycin resistance. For a and b, statistical significance between samples was calculated using an unpaired, two-tailed Student *t*-test; ns, no significant difference ($P > 0.05$). Data are shown as mean ± SD of 3 independent experiments.

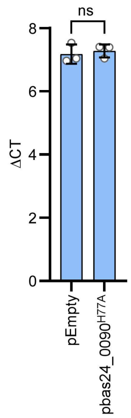

**Fig. S7. *bas24_0090* does not affect GAPS4$^{RiKo}$ expression.** GASP4 expression levels in the presence and absence of bas24_0090$^{H77A}$, as determined by quantitative real-time PCR (ΔCT). Statistical significance was calculated using an unpaired two-tailed Student *t*-test. ns, no significant difference ($P > 0.05$). Data are shown as mean ± SD of 3 independent experiments.

### Supplementary Tables

#### Table S1. Bacterial strains used in this study.

| Strain | Genotype | Source |
|---|---|---|
| *Escherichia coli* Neb5α | F$^-$ f80*lacZ*DM15D*(lacZYA-argF)* U169 *deoR recA1 endA1 hsdR17* ($r_K^-$, $m_k^+$) *gal* $^-$ *phoA supE44* l$^-$ *thi* $^-$*1 gyrA96 relA1* | NEB |
| *Escherichia coli* BW25113 | *lacI*$^q$, rrnBT14, D*lacZ*wJ16, *hsd*R514, D*ara*BAD AH33, D*rha*BAD LD78 | Lab collection |
| *Escherichia coli* BW25113 Δ*trxA*::kan | *lacI*$^q$, rrnBT14, D*lacZ*wJ16, *hsd*R514, D*ara*BAD AH33, D*rha*BAD LD79, *trxA* replaced with kanamycin resistance | Lab collection |
| *Escherichia coli* BW25113 Δ*trxA* | *lacI*$^q$, rrnBT14, D*lacZ*wJ16, *hsd*R514, D*ara*BAD AH33, D*rha*BAD LD79, *ΔtrxA* | Lab collection |
| *Escherichia coli* IYB5101 | *lacI*$^q$, rrnBT14, D*lacZ*wJ16, *hsd*R514, D*ara*BAD AH33, D*rha*BAD LD78, araB::T7RNAP-tetA | (*1*) |
| *Escherichia coli* TMB101 | *Escherichia coli* BW25113, araB::T7RNAP-tetA, λ-cI857 | This study |
| *Escherichia coli* DH5α λ-pir | F– φ80lacZΔ*M15* Δ(*lacZYA-argF*)*U169 recA1 endA1 hsdR17*(*rK– mK+*) *phoA supE44 λ– thi-1 gyrA96 relA1* Δ(*argF-lac*)169 pir+ | Lab collection |
| *Escherichia coli* MG1655 | F- lambda- ilvG- rfb-50 rph-1 | Lab collection |
| *Escherichia coli* K12 MG1655 ΔRM | *Escherichia coli* K-12 MG1655 Δ*mrr-hsdRMS-mcrBC* Δ*mcrA* = ΔRM with pBR322_ΔPtet F(pifA::zeoR) | Dr. Alexander Harms (ETH Zurich) |
| *Escherichia coli* BL21 (DE3) | *fhuA2 [lon] ompT gal (λ DE3) [dcm] ∆hsdS λ DE3 = λ sBamHIo ∆EcoRI-B int::(lacI::PlacUV5::T7 gene1) i21 ∆nin5* | Lab collection |
| *Escherichia coli* RiKo 2299/09 | Wild-type isolate | Prof. Ulrich Dobrindt (University of Münster) |
| *Vibrio parahaemolyticus* RIMD 2210633 | Wild-type clinical isolate | Prof. Kim Orth (University of Texas Southwestern Medical Center) |
| *Vibrio parahaemolyticus* T6SS1$^+$ effectorless | RIMD 2210633 Δ*hns*/Δ*tdhAS*/ Δ*vp_rs06130*/Δ*vp_rs06745*/*vp_rs06875*$^{AAA}$ | (*2*) |
| *Vibrio parahaemolyticus* T6SS1$^-$ effectorless | RIMD 2210633 Δ*hns*/Δ*tdhAS*/ Δ*vp_rs06130*/Δ*vp_rs06745*/ *vp_rs06875*$^{AAA}$/Δ*hcp1* | (*2*) |

**Table S2. Plasmids used in this study.**

| Plasmid | Origin of replication | Antibiotic resistance | Comments | Source |
|---|---|---|---|---|
| pBAD33.1 | ori15a | chloramphenicol | L-arabinose-inducible expression vector | Addgene #36267 |
| pGAPS4$^{MAVP-R}$ | ori15a | chloramphenicol | GAPS4 from *Vibrio parahaemolyticus* MAVP-R cloned in the pBAD33.1 MCS | (*3*) |
| pGAPS4$^{MAVP-R(D59A)}$ | ori15a | chloramphenicol | A D59A mutation introduced into pGAPS4$^{MAVP-R}$ | This study |
| pGAPS4a$^{MAVP-R}$ | ori15a | chloramphenicol | GAPS4a from *Vibrio parahaemolyticus* MAVP-R in the pBAD33.1 MCS | This study |
| pGAPS4b$^{MAVP-R}$ | ori15a | chloramphenicol | GAPS4b from *Vibrio parahaemolyticus* MAVP-R in the pBAD33.1 MCS | This study |
| pC-0_v5 | pSC101 | Ampicillin | A low-copy-number plasmid | Addgene #124425 |
| pGAPS4$^{RiKo}$ | pSC101 | Ampicillin | GAPS4 from *E. coli* RiKo 2299/09 cloned into pC-0_v5 along with 182 bp upstream containing the native promoter, and 235 bp downstream containing the native terminator sequences | This study |
| pGAPS4$^{RiKo(D49A)}$ | pSC101 | Ampicillin | A D49A mutation introduced into pGAPS4$^{RiKo(D49A)}$ | This study |
| pMBP-cas1 | pBR322 | Kanamycin | Cas1 cloned in T7-His-TEV-MBP plasmid | (*4*) |
| pMBP-GAPS4$^{MAVP-R}$ | pBR322 | Kanamycin | GAPS4$^{MAVP-R}$ cloned instead of Cas1 in the pMBP-cas1 plasmid, where GAPS4a is in-frame with the N-terminal MBP | This study |
| pBAD$^{K}$/Myc-His | pBR322 | Kanamycin | An arabinose-inducible expression vector | (*5*) |
| pbas24_0090 | pBR322 | Kanamycin | Bas24_0090 amplified from a Basel 24 phage and cloned into the pBAD$^{K}$/Myc-His MCS (untagged) | This study |
| pbas24_0090$^{H77A}$ | pBR322 | Kanamycin | A H77A mutation introduced into pbas24_0090 | This study |

| | | | | |
|---|---|---|---|---|
| pT7_3.8$^{H34A}$ | pBR322 | Kanamycin | T7_3.8 containing a H34A mutation cloned into pBAD$^{K}$/Myc-His MCS (untagged) | This study |
| pNF02-mChartreuse | Ori2 | Chloramphenicl | Constitutive expression of mChartreuse in *E. coli* | Addgene #219397 |
| pGFP | pBR322 | Kanamycin | mChartreuse, amplified from pNF02-mChartreuse, cloned into the pBAD$^{K}$/Myc-His MCS in-frame with the C-terminal Myc-6xHis tag of the pBAD$^{K}$/Myc-His plasmid | This study |
| pBAD$^{K}$/Myc-His-PoNe/i$^{Vp\ 12\text{-}\ 297/B}$ | pBR322 | Kanamycin | *Vibrio parahaemolyticus* 12-297/B VgrG1b with the downstream effector and immunity module PoNe/i, consisting of b5c30_rs14470-60 and the 3' gene, cloned in-frame with the C-terminal Myc-6xHis tag of the pBAD$^{K}$/Myc-His plasmid | (*6*) |
| pGEM-T | pBR322 | Ampicillin | A commercially available linear plasmid for TA cloning | Promega |
| pGEM$^{2.8OH}$-*trxA* | pBR322 | Ampicillin | *trxA-FLP* flanked by sequences homologous to those upstream and downstream of T7_2.8 was cloned into pGEM plasmid using TA cloning | This study |
| pGEM$^{3.8OH}$-*trxA* | pBR322 | Ampicillin | *trxA-FLP* flanked by sequences homologous to those upstream and downstream of T7_3.8 was cloned into pGEM plasmid using TA cloning | This study |
| pGEM$^{7.7OH}$-*trxA* | pBR322 | Ampicillin | *trxA-FLP* flanked by sequences homologous to those upstream and downstream of T7_7.7 was cloned into pGEM plasmid using TA cloning | This study |

| | | | | |
|---|---|---|---|---|
| pAC-ff-*FLP* | ori15a | chloramphenicol | A plasmid encoding FLP recombinase | Lab Collection |
| pIYPE69 | ori15a | Kanamycin | A plasmid containing a T7-packaging signal | (*7*) |

**Table S3. DNA primers used in this study.**

| Primer name | Sequence (5'-> 3')' | Use |
|---|---|---|
| TM573F | TTAAGAAGGAGATATACATATGTCAGGAGAGAAATCG | Amplify pGAPS4$^{MAVP-R}$ for site-directed mutagenesis to construct pGAPS4$^{MAVP-R(D59A)}$ |
| TM555R | CCTGAAATGTATGCATATTGTGCAGCAATACCATGCTGTTCTCGTTTTCC | |
| TM555F | GGAAAACGAGAACAGCATGGTATTGCTGCACAATATGCATACATTTCAGG | Amplify pGAPS4$^{MAVP-R}$ for site-directed mutagenesis to construct pGAPS4$^{MAVP-R(D59A)}$ |
| TM572R | CATCCGCCAAAACAGCCAAGCTT | |
| TM559R | ATCCGCCAAAACAGCCAAGCTTTTATTCATCGTTTAAGTTCCTAAAATTTGG | Amplify pGAPS4$^{MAVP-R}$ along with TM573F to construct pGAPS4a$^{MAVP-R}$ |
| TM560F | TTAAGAAGGAGATATACATATGAATAACGTTAGCACTTTCGTCC | Amplify pGAPS4$^{MAVP-R}$ along with TM572R to construct pGAPS4b$^{MAVP-R}$ |
| IY508R | TGATGTAGCCGTCAAGTTGTCATAA | Amplify the pC-o_V5 backbone to construct pGAPS4$^{RiKo}$ |
| IY139F | TGCTGCCACCGCTGAGCAAT | |
| TM712F | TGACAACTTGACGGCTACATCAGTTGTATTAGCTTACATATCTATGGCACAG | Amplify GAPS4$^{RiKo}$ from *E. coli* strain RiKo 2299/09 to construct pGAPS4$^{RiKo}$ |
| TM712R | TTATTGCTCAGCGGTGGCAGCATGTGCAGAGCATGAGACGCTATTTG | |
| AH33R | GAGTTGTAGACATACAAAGCAGCTATTCCATGTGTCTTTTTAGC | Amplify pGAPS4$^{RiKo}$ along with TM712F for site-directed mutagenesis to construct pGAPS4$^{RiKo(D49A)}$ |
| AH33F | GCTAAAAAGACACATGGAATAGCTGCTTTGTATGTCTACAACTC | Amplify pGAPS4$^{RiKo}$ along with TM712R for site-directed mutagenesis to construct pGAPS4$^{RiKo(D49A)}$ |
| TM224F | GATAACGGATCCGAATTCGAGCG | Amplify pMBP-*cas1* backbone to construct pMBP-GAPS4$^{MAVP-R}$ |
| TM224R | GGATTGGAAGTACAGGTTTTCCTCG | |
| TM604F | GGAAAACCTGTACTTCCAATCCATGTCAGGAGAGAAATCGAAGTCC | Amplify GAPS4$^{MAVP-R}$ from pGAPS4$^{MAVP-R}$ to construct pMBP-GAPS4$^{MAVP-R}$ |
| TM604R | GCTCGAATTCGGATCCGTTATCTTAGGACGAGCCAGGAGGAG | |
| TM377R | CATGGTTAATTCCTCCTGTTAG | Amplify pBAD$^{K}$/Myc-His backbone to construct pbas24_0090 |
| TM304F | AAGCTTGGGCCCGAACAAAAACTC | |
| TM733F | CTAACAGGAGGAATTAACCATGACTGAAGAATGGAAAAGAATACC | Amplify bas24-0090 from Basel 24 phage to construct pbas24_0090 |
| TM733R | GTTTTTGTTCGGGCCCAAGCTTTTAATATTCGTTATAAGTCTTTCTTAATACAATACAG | |

| | | |
|---|---|---|
| TM735R | ATTCAAAGGATTGCCATCAAGAGCTCGTGCATCTAGTTTCTCAT | Amplify pbas24_0090 along with TM733F for site-directed mutagenesis to construct pbas24_0090$^{H77A}$ |
| TM735F | TTATGAGAAACTAGATGCACGAGCTCTTGATGGCAATCCTTTGA | Amplify pbas24_0090 along with TM733R for site-directed mutagenesis to construct pbas24_0090$^{H77A}$ |
| TM787F | CTAACAGGAGGAATTAACCATGCGCAAGTCTTATAAACAATTC | Amplify T7_3.8 from T7 phage for site-directed mutagenesis to construct pT7_3.8$^{H34A}$ |
| TM788R | GTCTATATAATAACCTTTTGGTATAGGC | |
| TM787R | GTTTTTGTTCGGGCCCAAGCTTCTATCGGAATCGTGCGAATTG | Amplify T7_3.8 from T7 phage for site-directed mutagenesis to construct pT7_3.8$^{H34A}$ |
| TM788F | ACCAAAAGGTTATTATATAGACGCCATTGACGGCAATCCACTCAAC | |
| pBADfix_GIB_F_ (BJ) | AAGCTTGGGCCCGAACAAAAAC | Amplify pBADK/Myc-His backbone to construct pGFP |
| pBADfix_GIB_R_ (BJ) | CATGGTTAATTCCTCCTGTTAG | |
| BADfix_fluo_F_KZ | CTAACAGGAGGAATTAACCATGTCTAAAGGTGAAGAACTGTTCACC | Amplify mChartreuse from pNF02 to construct pGFP |
| fluo_myc_R_KZ | GTTCGGGCCCAAGCTTTTTGTAGAGCTCATCCATGCC | |
| TM738F | ACGAAGACGACGAAGAGTCCGAGGAAGCAGACGAAGACGGAGACTTCTAAGATCCGTCAGCCTGCAGTTC | Amplify trxA-FLP flanked by sequences homologous to those upstream and downstream of T7 gene 2.8 and cloned in pGEM-T vector using TA cloning to construct pGEM$^{2.8OH}$-trxA |
| TM738R | TTCCTTTAGCGCCGTAACCTGCCATGTCGCCTCCCTTTGCGTATATCACAGTGTAGGCTGGAGCTGCTTC | |
| TM739F | CTAAAGGGAGACCACAGCGGTTTCCCTTTGTTCGCATTGGAGGTCAAATAGATCCGTCAGCCTGCAGTTC | Amplify trxA-FLP flanked by sequences homologous to those upstream and downstream of T7 gene 3.8 and cloned in pGEM-T vector using TA cloning to construct pGEM$^{3.8OH}$-trxA |
| TM739R | TATCGGAATCGTGCGAATTGTCCATGCAATTCCCTCCTAGTTCTATAAATGTGTAGGCTGGAGCTGCTTC | |
| TM741F | TGTAGCCCGTAGCTCCGGTGGCGGTATCAACATTTAATCAGGAGGTTATCGATCCGTCAGCCTGCAGTTC | Amplify trxA-FLP flanked by sequences homologous to those upstream and downstream of T7 gene 7.7 and cloned in pGEM-T vector using TA cloning to construct pGEM$^{7.7OH}$-trxA |
| TM741R | CAAGTCCTGTTCGTTTCTCAGCCATTAAATGTGTCTCCATGTCTTACGCTGTGTAGGCTGGAGCTGCTTC | |
| 105F | ATTATCGGTGGTGGCTCTAA | Amplify the regions flanking T7-gp 2.8 (used for phage competition assay to distinguish wild-type from mutant T7 phage) |
| SM24R8 | ATTACTGCAGTTAGCGCCGTAACCTGCCAT | |
| TM769F | AGACTTGGCACACACCATTG | Amplify GAPS4$^{RiKo}$ for RT-PCR |
| TM769R | GCTCTTGCGTTATCAACGAG | |

| | | |
|---|---|---|
| TM110F | CGTCAGCTCGTGTTGTGAAA | Amplify *E. coli* 16s rRNA for RT-PCR |
| TM110R | TGTGTAGCCCTGGTCGTAAG | |
| TM623F | GACGGATTCATCGTTGGGGTC | Amplify λ-cI857 phage genome for RT-PCR |
| TM634R | CAGCTGAATGGTGCAGTTCTG | |

**Table S4. List of phages used in this study.**

| Phage | Family | Source |
|---|---|---|
| T4 | *Myoviridae* | Lab collection |
| T5 | *Siphoviridae* | Lab collection |
| T7 | *Podoviridae* | Lab collection |
| λ-vir | *Siphoviridae* | Lab collection |
| P1-vir | *Myoviridae* | Lab collection |
| Basel 01 | *Drexlerviridae; Braunvirinae* | (*8*) |
| Basel 02 | *Drexlerviridae; Braunvirinae* | |
| Basel 03 | *Drexlerviridae; Braunvirinae* | |
| Basel 04 | *Drexlerviridae; Tempevirinae* | |
| Basel 05 | *Drexlerviridae; Tempevirinae* | |
| Basel 06 | *Drexlerviridae; Tempevirinae* | |
| Basel 07 | *Drexlerviridae; Tempevirinae* | |
| Basel 08 | *Drexlerviridae; Tempevirinae* | |
| Basel 09 | *Drexlerviridae; Tempevirinae* | |
| Basel 10 | *Drexlerviridae; Tempevirinae* | |
| Basel 11 | *Drexlerviridae; Tempevirinae* | |
| Basel 12 | *Drexlerviridae; Tunavirinae* | |
| Basel 13 | *Drexlerviridae; Tunavirinae* | |
| Basel 14 | *Siphoviridae* | |
| Basel 15 | *Siphoviridae* | |
| Basel 16 | *Siphoviridae* | |
| Basel 17 | *Siphoviridae* | |
| Basel 18 | *Siphoviridae* | |
| Basel 20 | *Siphoviridae, Queuovirinae* | |
| Basel 21 | *Siphoviridae, Queuovirinae* | |
| Basel 22 | *Siphoviridae, Queuovirinae* | |
| Basel 23 | *Siphoviridae, Queuovirinae* | |
| Basel 24 | *Siphoviridae, Queuovirinae* | |
| Basel 25 | *Siphoviridae, Queuovirinae* | |
| Basel 26 | *Demerecviridae; Markadamsvirinae* | |
| Basel 27 | *Demerecviridae; Markadamsvirinae* | |
| Basel 28 | *Demerecviridae; Markadamsvirinae* | |
| Basel 29 | *Demerecviridae; Markadamsvirinae* | |

| | |
|---|---|
| Basel 30 | *Demerecviridae; Markadamsvirinae* |
| Basel 31 | *Demerecviridae; Markadamsvirinae* |
| Basel 32 | *Demerecviridae; Markadamsvirinae* |
| Basel 33 | *Demerecviridae; Markadamsvirinae* |
| Basel 34 | *Demerecviridae; Markadamsvirinae* |
| Basel 35 | *Myoviridae; Tevenvirinae* |
| Basel 36 | *Myoviridae; Tevenvirinae* |
| Basel 37 | *Myoviridae; Tevenvirinae* |
| Basel 38 | *Myoviridae; Tevenvirinae* |
| Basel 39 | *Myoviridae; Tevenvirinae* |
| Basel 40 | *Myoviridae; Tevenvirinae* |
| Basel 41 | *Myoviridae; Tevenvirinae* |
| Basel 42 | *Myoviridae; Tevenvirinae* |
| Basel 43 | *Myoviridae; Tevenvirinae* |
| Basel 44 | *Myoviridae; Tevenvirinae* |
| Basel 45 | *Myoviridae; Tevenvirinae* |
| Basel 46 | *Myoviridae; Tevenvirinae* |
| Basel 47 | *Myoviridae; Tevenvirinae* |
| Basel 48 | *Myoviridae; Vequintavirinae* |
| Basel 49 | *Myoviridae; Vequintavirinae* |
| Basel 50 | *Myoviridae; Vequintavirinae* |
| Basel 51 | *Myoviridae; Vequintavirinae* |
| Basel 52 | *Myoviridae; Vequintavirinae* |
| Basel 53 | *Myoviridae; Vequintavirinae* |
| Basel 54 | *Myoviridae; Vequintavirinae* |
| Basel 55 | *Myoviridae; Vequintavirinae* |
| Basel 56 | *Myoviridae; Vequintavirinae* |
| Basel 57 | *Myoviridae; Vequintavirinae* |
| Basel 58 | *Myoviridae; Vequintavirinae* |
| Basel 59 | *Myoviridae; Vequintavirinae* |
| Basel 60 | *Myoviridae; unclassified* |
| Basel 61 | *Myoviridae; unclassified* |
| Basel 62 | *Myoviridae; unclassified* |
| Basel 63 | *Myoviridae; Ounavirinae* |

| | | |
|---|---|---|
| Basel 64 | *Autographiviridae; Studiervirinae* | |
| Basel 65 | *Autographiviridae; Studiervirinae* | |
| Basel 66 | *Autographiviridae; Studiervirinae* | |
| Basel 67 | *Autographiviridae; Studiervirinae* | |
| Basel 68 | *Autographiviridae; Studiervirinae* | |
| Basel 69 | *Schitoviridae; Enquatrovirinae* | |
| λ-CI857 | *Siphoviridae* | (*9*) |
| T7Δ2.8::trxA-FLP | *Podoviridae* | This study |
| T7Δ2.8 | *Podoviridae* | This study |
| T7Δ2.8Δ7.7::trxA-FLP | *Podoviridae* | This study |
| T7Δ2.8Δ7.7 | *Podoviridae* | This study |
| T7Δ2.8Δ7.7Δ3.8::trxA-FLP (T7Δ3HE) | *Podoviridae* | This study |

### Supplementary Datasets

**Dataset S1. A list of GAPS4 homologs identified in this study**

### Supplementary Files

**File S1. The concatenated amino acid sequences of GAPS4a and GAPS4b homologs identified in this study**

**File S2. AlphaFold3 structure prediction for $GAPS4^{MAVP-R}$**

**File S3. AlphaFold3 structure prediction for $GAPS4^{RiKo}$**
